# An Epigenetic Signature of Vulnerable Neurons is Under Selective Pressure Associated with Longevity Across Placental Mammals

**DOI:** 10.64898/2026.08.20.745782

**Authors:** Ghada Abdelhady, Qiao Su, Andrew Z. Wang, Rajee Ganesan, BaDoi N. Phan, Heather H. Sestili, Vijay Cherupally, The Vertebrate Genomes Project Consortium Phase I, Andreas R. Pfenning

**Author notes:** **CORRESPONDING AUTHOR:** ARP.

## Abstract

Age is the primary risk factor for neurodegenerative diseases, which are characterized by cell-type-specific vulnerability ^1^. Yet brain-aging mechanisms remain unclear given the complex, interacting age-associated pathways across diverse neural cell types. Here, we dissect cell type- cell state-specific aging gene regulatory programs and their contribution to cellular vulnerability by leveraging epigenomics, AI methodology, and natural lifespan diversity across placental mammals. Applying the TACIT method, we associated lifespans of 240 placental mammals to the predicted open chromatin levels of over 3 million orthologous loci across 18 cortical cell types. We identified thousands of lifespan-associated open chromatin regions, enriched near genes associated with hallmarks of aging, which stratified greatly by cell type. For example, regions near mitochondrial genes showed differential selective pressure in long-lived species in energetically-demanding layer V ET neurons, while regions near inflammatory response genes were under selective pressure in glial populations. We next asked whether regions linked to vulnerable or resilient neurons in the human brain were under differential selective pressure in longer lived species. Using an adaptive representation learning approach, we decompose intrinsic aging programs from systemic effects in the prefrontal cortex and define an aging signature predictive of cell-type-specific vulnerability. In Alzheimer’s disease, this intrinsic aging signature more strongly predicts vulnerability than systemic effects. Active regions in vulnerable neurons showed lower predicted activity in species with longer lifespans, suggesting selective pressure to down-regulate the vulnerability-associated networks. Overall, our findings argue against a single master regulator of aging, instead implicating different hallmarks across different cell types.

## INTRODUCTION

Age is the primary risk factor for neurodegenerative diseases, which are characterized by progressive loss of neurons, with cell type-specific selective vulnerability being a key feature ^1^. Yet, the causal mechanisms underlying this selective loss are unclear. Aging and neurodegeneration have been associated with a variety of molecular hallmarks, including DNA damage, epigenetic alterations, inflammation, oxidative phosphorylation, senescence, and protein misfolding, and age-related synaptic alterations ^2–4^. This positions aging as a route for tracing the causal drivers for neuronal loss in neurodegenerative diseases.

Identification of shared and distinct molecular mechanisms between normal aging and neurodegeneration can provide a more profound understanding of the initiating events that drive the disease onset. With the goal of maximizing the period of life spent free from disease, a key question is: which early molecular programs of brain aging predict and potentially drive neuronal vulnerability to degeneration? Addressing this requires delineating trajectories that characterize normal aging. And here it’s critical, to choose the reference scale for evaluating normal aging.

Although human longevity is heritable ^5^, confounding factors and the limited genetic variation across the human populations limits the statistical power of human genetics alone to identify factors driving aging in specific cell types and contexts. An opportunity lies in leveraging the variability in longevity across mammals. While this variation in lifespan is partly influenced by ecological factors such as social structure, mobility and nutrition, it is also driven by longevity adaptability through gene regulation, adaptive responses, and stress resilience ^6^. Within humans, aging reflects a species-specific form of adaptability, with context-dependent regulatory changes giving rise to heterogeneous aging trajectories across individuals ^7,8^. At the cellular level, aging trajectories can further diverge within an individual, indicating variable adaptability and susceptibility to aging and age-related neurodegeneration. Across these hierarchical scales –from species to individuals to cells– a unifying theme is the capacity to adapt over time. However, whether these adaptive processes are shared, causally linked, or operate independently across scales remains unclear (Figure 1, top panel).

**Figure 1.**
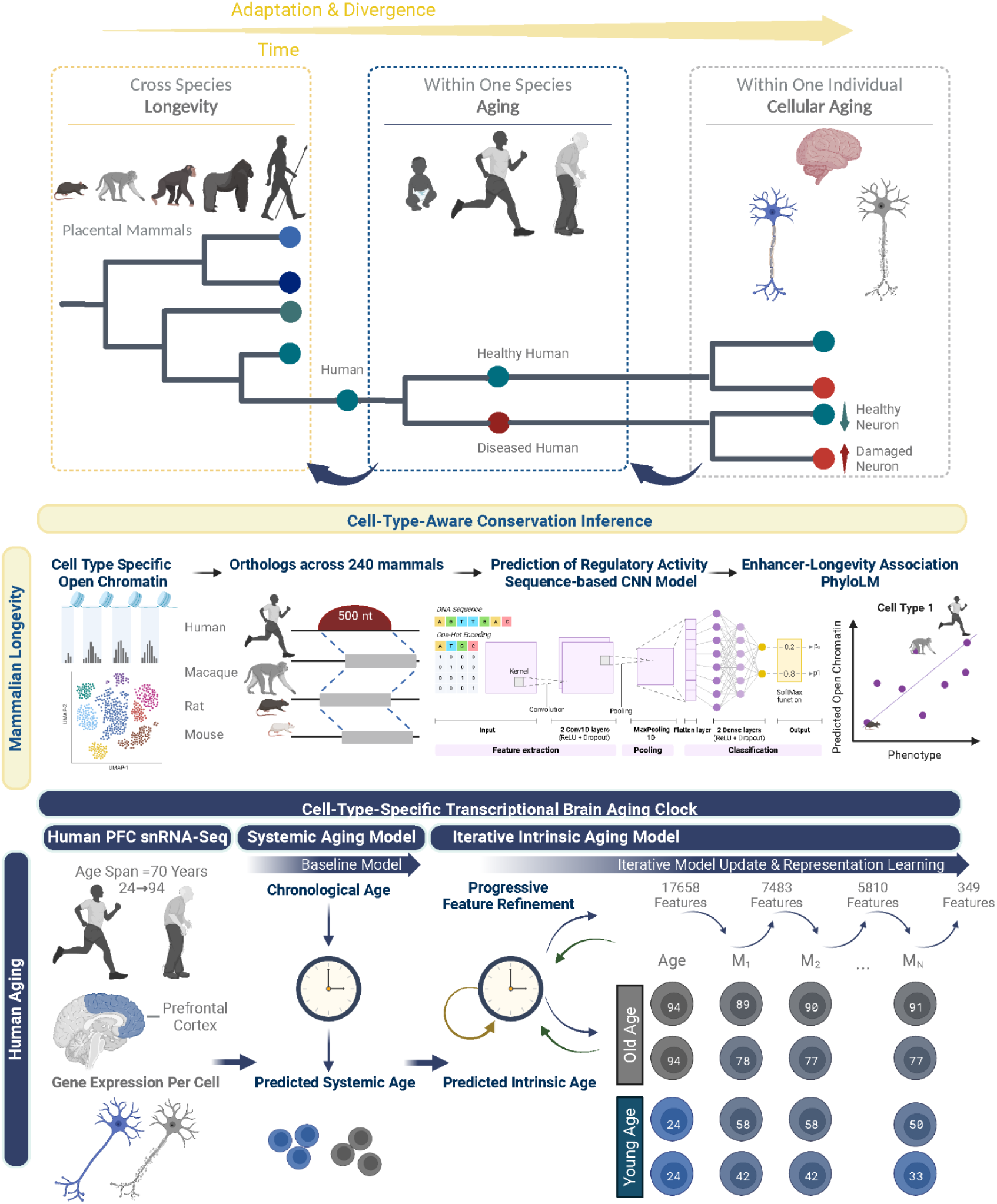
Hierarchical Adaptation and Divergence. Overview of hierarchical adaptation across scales from evolutionary differences in longevity across species to organismal aging in humans, with a focus on cell-type-specific aging programs (top panel). Cell-Type-Aware longevity evolution framework spanning 240 mammals (middle panel). Adaptive Feedback Model of the Transcriptional Brain Aging Clock: Architecture of the iterative elastic regression model (iENR) demonstrating refinement of critical features for accurate prediction of cellular aging (bottom panel). Schematics are created in BioRender.com.

In this study, we investigated the selective pressures associated with longevity evolution across cortical cell types, including neurons, glia, and endothelial cells. Using an atlas of 240 mammalian genomes^9^, we applied the Tissue-Aware Conservation Inference Toolkit (TACIT)^10^ to identify open chromatin regions (OCRs) whose evolutionary divergence tracked divergence in species lifespan.

Focusing on the adaptive processes in the brain and particularly in post-mitotic neurons, which provide a unique context in which identity and functional specialization must be preserved through tightly controlled transcriptional programs across the lifespan of an individual ^11,12^. The functional activity of a neuron is a product of systemic factors –reflective of its environment and connectivity with other neurons, and intrinsic factors –reflective of the cell’s identity and its unique gene regulatory architecture. As such, functional constraints are not only imposed on the evolution of neural circuitry for an organism’s survival^13^ but it’s expected to be imposed on the neuronal identity as well.

Understanding how intrinsic identity programs and systemic inputs jointly shape neuronal aging requires quantitative frameworks capable of capturing molecular age across scales. One approach to studying the heterogeneity in cellular aging is through molecular aging clocks: machine learning models trained to predict the age of a sample based on biological features. The first epigenetic aging clocks used bulk-tissue DNA methylation levels at CpG sites as biomarkers of age-related epigenetic drift in humans^14,15^. Recently, a pan-mammalian aging clock spanning 185 species identified conserved methylation changes across mammals, suggesting that aging is evolutionarily conserved ^16^. Such conservation points to a shared epigenomic program of aging, in which dynamic changes in DNA methylation contribute to systematic shifts in the transcriptome. Consistent with this, stress-related DNA methylation changes arising from environmental exposure can alter gene expression ^15,17^.

The transcriptome provides a more interpretable and functionally testable readout of aging-associated changes. However, transcriptomic aging clocks have faced challenges related to technical noise and limited sample sizes, reducing reproducibility ^18^. Moreover, most approaches have relied on bulk-tissue data, which fails to capture cellular aging heterogeneity. Recent efforts using ElasticNet models with single-nucleus RNA sequencing (snRNA-Seq) reported that ElasticNet outperformed other models in terms of predictive performance, interpretability, and efficiency. However, these approaches often rely on binarized expression of a subset of highly variable genes and were trained on model organisms ^19^, limiting its ability to capture the full spectrum of gene expression dynamics and identify novel human-specific biomarkers.

These limitations underscore the need for interpretable, single-cell transcriptomic aging clocks capable of resolving genome-wide, cell-type-specific transcriptional aging signatures in the human brain. While traditional differential gene expression analyses can identify genes with significant mean changes, they rely on discrete group comparisons. In contrast, transcriptional aging clocks model aging as a continuous process, amplifying sensitivity to gradual continuous transcriptional shifts across cells. For instance, a gene may not be individually differentially expressed but jointly with other genes, it’s crucial for encoding the age of a cell. Transcriptional aging clocks leverage cellular heterogeneity in gene expression as an informative signal for prediction, rather than treating it as noise, revealing aging-associated transcriptional programs that are obscured in group-averaged differential expression analyses.

Here, we present an adaptive, ElasticNet-based iterative learning framework to resolve aging at the single-cell resolution. Using human prefrontal cortex snRNA-Seq data^20^, we develop a cell-type-specific transcriptomic aging clock, which unlike standard transcriptomic clocks, it decomposes shared and cell-specific aging signals through iterative refinement. Through this framework, we identified an intrinsic aging signature that is orthogonal to chronological aging signals. Cell-intrinsic age was associated with mitochondrial-oxidative phosphorylation as key cell-autonomous processes driving molecular aging. We find that neuronal populations enriched for gene expression programs positively associated with intrinsic aging exhibit relative resilience, maintaining adaptive transcriptional responses with age. In contrast, neuronal populations enriched for negatively associated programs are selectively vulnerable, potentially reflecting failure to engage protective mechanisms. As aging advances and neurodegenerative risk rises, accumulating cellular insults disrupt neuroprotective programs and whether a cell is equipped to adapt to these perturbations ultimately shapes its aging trajectory.

To further investigate the causal contribution of aging-associated regulatory elements to cellular vulnerability, we intersected human aging with the evolution of longevity across mammals and at higher scale, across vertebrates. By distinguishing conserved from divergent aging pathways, this approach enables the identification of regulatory mechanisms that may contribute to healthy aging. We show that these age-associated evolutionary signals can be used to identify vulnerable neuronal states. More broadly, our results suggest that cell type- and cell state-specific regulatory networks in humans are associated with related evolutionary pressures across placental mammals.

## RESULTS

### A multi-scale framework integrating a transcriptional brain aging clock with evolutionary longevity

To delineate shared and species-specific aging pathways across placental mammals and identify regulatory mechanisms that may promote healthy aging, we designed a multi-scale framework to investigate cell-type-specific aging programs and their contribution to cellular vulnerability in neurodegenerative disease. First, our approach connects **cross-species lifespan** measurements with aging within the human population. Second, our approach separates out **intrinsic factors**, which are likely to be cell autonomous effects, versus **systemic factors that** are more likely to be influencing cells broadly across the entire tissue (Figure 1).

Our approach integrates these scales through seven components: (1) resolving neuronal and non-neuronal cortical cell-type-specific longevity-associated regulatory elements (LARs) across 240 mammals; (2) development of a transcriptional brain aging clock to disentangle systemic aging from intrinsic aging in cortical neurons within the human population; (3) resolving cell-type-specific intrinsic aging; (4) linking intrinsic age to cellular vulnerability and aging hallmarks; (5) identification of differentially accessible regulatory elements in vulnerable cell types; (6) integration of cross-species and human-focused analyses to identify shared regulatory elements connecting mammalian longevity with human aging at the cell-type level, vulnerability-longevity-associated regulatory elements (VLRs) and (7) investigation of evolutionary conservation of enhancer-gene synteny for VLRs across 577 vertebrate species. Unlike prior approaches that examine aging at a single scale or modality, this framework systematically integrates evolutionary, tissue-level, and cell-intrinsic dimensions to resolve how regulatory programs of aging emerge and contribute to disease vulnerability.

### Longevity Selective Pressure Across Placental Mammals Resolved At the Cell Type Level

Aging is a heterogeneous process shaped by multiple interconnected hallmarks, including genomic instability, telomere attrition, epigenetic alterations, loss of proteostasis, mitochondrial dysfunction, cellular senescence, inflammation, and altered intercellular communication ^3,4^. However, as previously reported, dissecting the interconnectedness among these hallmarks and determining their relative contributions to aging remains a challenge ^4^. Here, we address this from a cell-type-specific perspective. As cellular properties progressively change with age ^21^, cellular trajectories can diverge even within the same individual. This variability suggests that distinct cell types encode different layers of aging-related information and therefore exhibit differential susceptibilities and adaptive capacities during aging. Accordingly, we resolve aging hallmarks in a cell-type-specific context, revealing that the molecular programs underlying longevity are not uniform, but instead differ across cellular identities.

We leveraged genomic data from 240 species of placental mammals available through the Zoonomia consortium^9^ to investigate associations between cell type-specific open chromatin and longevity using the TACIT method. Open chromatin status was inferred across mammals by first identifying cell-type-specific open chromatin regions from single-nucleus ATAC-seq (snATAC-seq) data generated in four species: humans^22,23^, rhesus macaques^24^, rats^25,26^ and mice^27–29^ (Methods). Convolutional Neural Networks (CNNs) were trained to learn the sequence features underlying open chromatin levels across 18 cortical neural cell types and neuron subtypes. For almost all cell types, our models were able to predict cell type-specific open chromatin signatures on held out orthologous regions (average AUROC = 0.73 and AUPRC = 0.7). We then applied the cell type-specific CNNs to predict chromatin accessibility for orthologous regions across other mammalian genomes (Methods, Extended Data Figure 1A). As a result, for each of the **18 cell types**, we obtained predictions for **3,180,197 open chromatin regions** across **240 species**. To assess biological validity, we evaluated models trained on human OCRs by predicting enhancer activity in orthologous regions across 240 mammalian genomes for each cortical cell type. As expected for human OCRs, predicted accessibility declined progressively with increasing evolutionary distance from humans (Extended Data Figure 1B). This decline reflects the evolutionary decay of enhancer activity, accessibility becoming less conserved in more distant lineages.

We next associated the predicted open chromatin levels to longevity phenotypes. To isolate longevity from the confounding effects of body size, we annotated the longevity quotient (LQ), which controls for the relationship between lifespan and body size ^30–32^ (Figure 1A). We then applied a phylogenetic linear model to associate LQ with predicted open chromatin while accounting for phylogenetic relatedness among species. We found an average of 11,379 OCRs positively-associated with longevity and 15,151 OCRs negatively-associated with longevity, suggesting substantial evolutionary pressure across cell types for species longevity (Supplementary Tables 1-2). The most human-specific longevity-associated peak (hg38:chr12:6,284,924–6,285,425) was identified in microglia, showing the largest gain in predicted chromatin accessibility in humans relative to 28 non-human primates and a longevity-association coefficient of 2.9 (FDR-adjusted p = 3.46 × 10⁻⁴). This peak was located near *PLEKHG6*, a RhoA-activating guanine nucleotide exchange factor whose primate-specific isoform supports brain development ^33^ (Extended Data Figure 2A-B). We return to microglia below, where cell-type-resolved analysis reveals their role in longevity and disease risk.

To uncover the biological processes underlying longevity across species, we tested the enrichment of the complete set of LARs independent of cell-type identity. Pathways related to DNA damage response, apoptosis, and regulation of mitophagy were positively associated with longevity. These findings are consistent with previous reports in bats, where enhanced DNA repair and autophagy mechanisms contributed to extended lifespan ^34^. In contrast, processes related to histone acetylation and deacetylation were negatively associated with longevity (Extended Data Figure 2C), suggesting that disruption of chromatin organization and epigenetic regulation contribute to reduced lifespan. Collectively, these patterns align with previously described higher-order hallmarks of aging ^4^.

Resolving LARs at the cell-type level revealed substantial heterogeneity in the biological programs associated with lifespan, indicating that aging hallmarks manifest differently across cellular identities (Figure 2B). Among excitatory neurons, L2.3.IT and L4.5.IT exhibited negative associations with pathways regulating intrinsic apoptotic signaling in response to DNA damage. In contrast, L5.ET neurons showed positive associations with responses to topologically incorrect proteins and regulation of synaptic plasticity, while displaying negative associations with mitochondrial membrane permeability, apoptotic mitochondrial changes, and histone H4 acetylation. Interestingly, positive regulation of calcium exocytosis was negatively associated in L2.3.IT, L4.5.IT and L5.ET. Inhibitory interneuron populations exhibited distinct longevity-associated signatures. PVALB interneurons showed positive associations with calcium-dependent exocytosis, consistent with maintenance of fast-spiking synaptic transmission. SST interneurons were positively enriched for regulation of neuronal synaptic plasticity, whereas VIP interneurons were positively associated with histone H4 acetylation marks, including H4-K5 and H4-K8 acetylation.

**Figure 2.**
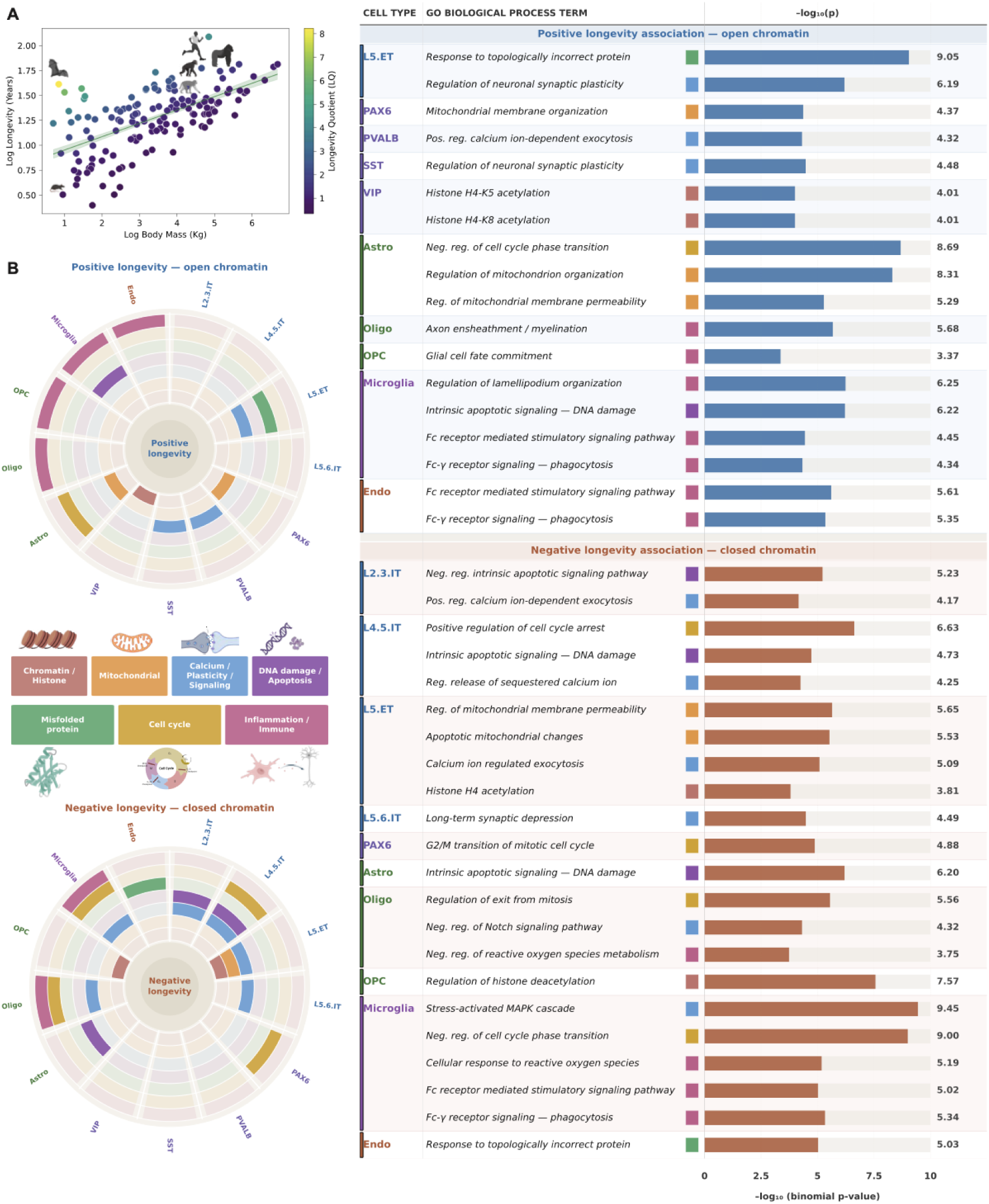
Cell-Type Resolved Longevity Hallmarks. A. Log Longevity regressed on Log Body Mass for mammals from the Zoonomia^9^ Database. Each point represents one species colored based on the gradient of the longevity quotient. B. Gene ontology enrichment analysis for cell-type-specific chromatin regions associated with longevity (positive=top, negative=bottom). Background was set to the whole genome using GREAT. Pathways were grouped into seven major biological categories: (1) Chromatin/Histone, (2) Mitochondrial, (3) Calcium/Synaptic signaling and plasticity, (4) DNA damage/Apoptosis, (5) Misfolded proteins, (6) Cell cycle, (7) Inflammation/Immune (left). Detailed pathways for each cell type with level of significance are shown (right).

Glial populations were enriched for pathways involved in inflammation and immune regulation. Astrocytes exhibited positive associations with mitochondrial organization, and regulation of mitochondrial membrane permeability, alongside negative association with DNA damage-induced apoptotic signaling. Microglia displayed positive enrichment for lamellipodium organization, apoptotic signaling in response to DNA damage, and Fc receptor-mediated phagocytic signaling pathways, highlighting immune surveillance and phagocytic activity as longevity-associated features. However, these same immune-related pathways, including Fc receptor signaling, stress-activated MAPK cascades and response to reactive oxygen species, were negatively associated in other contexts, indicating a complex balance between protective immune activation and chronic inflammatory signaling during aging, suggesting different microglial states. As expected, oligodendrocytes were positively associated with axon ensheathment and myelination, confirming the preservation of myelin maintenance programs in longer-lived species, while negatively associated with mitotic exit, Notch signaling repression, and reactive oxygen species metabolic regulation. OPCs were positively associated with glial cell fate commitment and negatively associated with histone deacetylation.

Furthermore, we observed that negatively LARs have a higher proportion in most cell types relative to the positively LARs. Notable exceptions included parvalbumin (PV) interneurons, glial populations, and endothelial cells, which exhibited approximately equal proportions of negative and positive LARs (Extended Data Figure 2D).

To assess the disease relevance of LARs, we examined their overlap with Alzheimer’s disease (AD) genetic risk loci. This analysis identified a microglia-specific negatively LAR located near the AD risk gene *BIN1* that overlapped four AD-associated single nucleotide polymorphisms (SNPs) (Extended Data Figure 2E and 3A-C). This pointed to microglial regulatory variation as a shared substrate linking longevity and AD risk. Thus, our framework can prioritize cell-type-specific regulatory elements that are both evolutionarily associated with longevity and enriched for disease-relevant genetic variation in a cell type highly implicated with AD pathogenesis.

**Figure 3.**
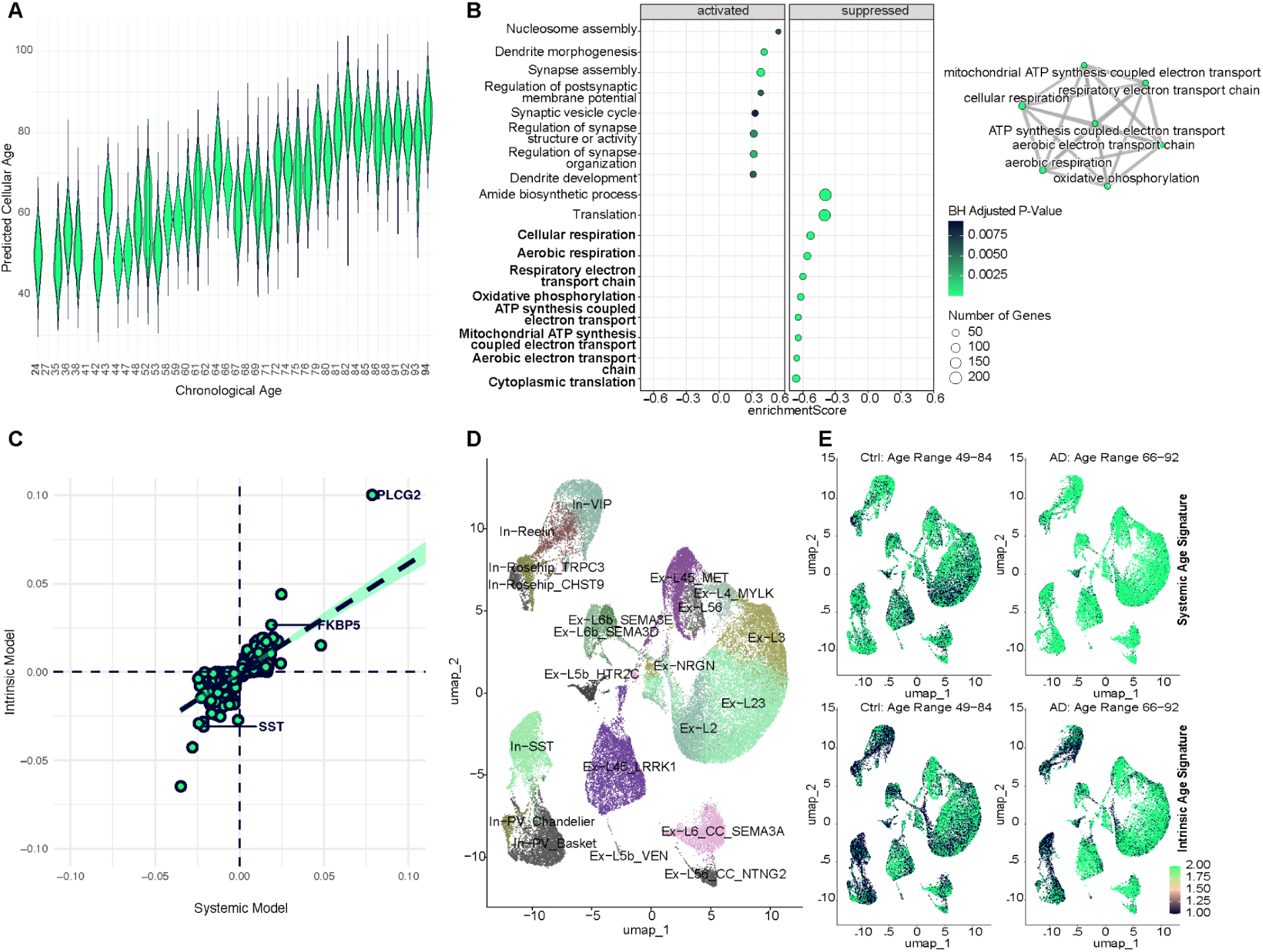
Utility of iterative elastic net regression for cellular age prediction, feature selection and profiling cellular heterogeneity. A. Violin plots of predicted cellular age (y-axis) based on each subject’s chronological age (x-axis) after the first iteration of model training, highlighting distribution of cellular age estimates across subjects with age range 24-94 years. B. Gene set enrichment analysis (GSEA) dot plot and enrichment map for genes with stronger association with intrinsic age relative to age, ranked by the difference between intrinsic age coefficient and age coefficient. C. Scatterplot of feature coefficients after model refinement, showing the first iteration (x-axis; systemic model) versus the final iteration (y-axis; intrinsic model). Genes with increased absolute weight in the final iteration, including *PLCG2, FKBP5 and SST*, are highlighted. D. Low dimensionality projection of cortical neuronal cell types from Anderson et al.^36^ dataset comprising AD patients and age-matched controls with age range 49-92 years. E. Low dimensionality projection feature plot of systemic age signature score (top) and intrinsic age signature score (bottom) in AD patients and age-matched controls. Color scale is uniform across all plots by scaling plots for all features and conditions to the maximum expression value for the feature with the highest overall expression.

Among neuronal populations, positive LARs in In-PV interneurons overlapped the largest number of AD-associated SNPs, consistent with the uncommonly high proportion of positive LARs observed in this cell type relative to other neuronal populations (Extended Data Figure 2E). Together, these findings highlight candidate regulatory loci that may contribute to selective neuronal vulnerability during aging and neurodegeneration, motivating a focused investigation of neuronal cell-type-specific aging trajectories in the human brain.

### Predicting neuronal age across and within human subjects using snRNA-Seq from prefrontal cortex post-mortem tissue

The cell-type specificity of longevity-associated selection implies that the programs governing neuronal vulnerability and resilience are themselves under selective pressure. For example, OCRs associated with neuronal vulnerability may be selected to have lower accessibility in longer-lived species, whereas OCRs associated with resilience may be selected for higher accessibility. Interpreting such a signal, however, requires decomposing not only chromatin accessibility but the downstream gene expression profile of a cell, which is a product of both intrinsic factors reflective of the cell’s identity or current state, and systemic factors imposed by the tissue environment surrounding the cell. Disentangling these influences is essential for understanding cell-autonomous aging and vulnerability, particularly in neurodegenerative diseases.

To disentangle these influences in the human brain, we turned from cross-species chromatin accessibility to single-nucleus transcriptomes of post-mortem prefrontal cortex. Our approach is an iterative elastic net regression model (iENR) that begins with a standard ElasticNet predictor of chronological age, capturing dominant systemic aging effects. The model then iteratively refines age predictions by using outputs from previous iterations as targets, progressively enriching for gene programs that explain cell-specific deviations from the shared aging trajectory (Supplementary Tables 3-4). The adaptive feedback-based aging clock enabled us to disentangle systemic aging uniformly affecting large populations of cells across the tissue from cell-intrinsic aging, in which more cell-autonomous mechanisms influence its molecular age. Gradual adaptation allowed for the identification of cells at different levels of aptitude to molecular aging within each human subject (age range 24-94 years) (Figure 3A), suggesting differential vulnerability to aging.

To gain a better understanding of the mechanisms underlying brain aging, we regressed the full set of 17,658 features on chronological age using ordinary least squares linear regression model (OLS), while accounting for the mean number of transcripts counts per subject. We, then, employed Gene Set Enrichment Analysis (GSEA) to the set of features significantly associated with age (positively or negatively) ranked by their age coefficient (Extended Data Figure 5A). As expected for the systemic aging-associated signature, a module of related terms included DNA damage response, chromatin organization and remodeling was consistent with the enrichment of LARs across mammals, independent of cell-type identity (Extended Data Figure 2C). Although this demonstrates the ability of our single-cell analysis to capture genes tied to known hallmarks of aging, the model may not capture the latent mechanisms driving heterogeneity in cellular aging within an individual.

To address this, we trained a second OLS model by regressing features on intrinsic age, identifying genes more strongly associated with intrinsic age than with chronological age, ranked by the difference between the intrinsic-age and chronological-age coefficients. Notably, nuclear-encoded genes showed a stronger negative association with intrinsic age and were enriched for cellular energy metabolism pathways dependent on mitochondrial ATP-synthesis-coupled electron transport and oxidative phosphorylation (OXPHOS) (Figure 3B, Extended Data Figure 5B). In other words, neurons with lower intrinsic age (molecularly younger) exhibited relatively higher expression of nuclear-encoded OXPHOS pathways.

To note, the iENR and OLS models are complementary but distinct: iENR model defines a gene’s membership in the intrinsic aging signature, whereas the OLS model quantifies the strength of each gene’s association with predicted intrinsic age versus chronological age. As expected, the iENR-defined intrinsic aging signature was a subset of the genes associated with intrinsic age in the OLS model.

### Modeling cellular heterogeneity in cortical neuronal cell types in healthy aging and disease

Following the confirmation of the utility of the model’s solution to separate intrinsic and systemic signatures of aging, we next sought to interpret the disparity in the implication of genes to the aging process at sequential resolutions of the model. For instance, we identified patterns for how the coefficient for a group of genes shifted with each model’s iteration. Some genes were only associated with the target variable in the first iteration and then their coefficient dropped to zero in the next iteration, indicating that systemic effects during aging are driving the expression of those genes. Another set of genes’ coefficients steadily increased or decreased while maintaining the directionality, and a few genes kept changing directionality in association with age (Figure 3C). The gene most strongly contributing to the intrinsic aging signature in the positive direction was *PLCG2*, which controls mitochondrial respiration and has been linked with vulnerability and resilience in Alzheimer’s disease ^35^. Another interesting gene is FK506-binding protein 51 (*FKBP5/FKBP51*), stress response modulator, exhibited a steady increase in its coefficient and maintained a positive directionality in association with intrinsic age. Its expression was selective to excitatory neurons, particularly L2, L2-3, L5-6 and L6 excitatory neurons and its expression increased with age (Extended Data Figure 5D, Top). In the opposite direction, XKR4 (XK Related 4) maintained a negative directionality in association with intrinsic age. XKR4 is involved in phosphatidylserine exposure on the cell surface for elimination of unwanted synapses ^36^. Its negative association with age could be contributing to the decline in synapse number with aging ^37^. Somatostatin (SST) is another interesting gene that was negatively associated with intrinsic age. As anticipated, *SST* expression was primarily localized to In-SST and, to a lesser extent, in In-Reelin, showing a decline with age (Extended Data Figure 5D, Bottom), consistent with previous findings. This underscores the model’s strong intrinsic cell-type-specific signal-to-noise ratio, even for rare populations. The adaptive refinement of genes from a broad set to a specific important subset supports the use of our iterative approach for disentangling systemic and intrinsic gene signatures.

Given the heterogeneous nature of aging, focusing on single genes is not sufficient nor comprehensive for capturing cell-autonomous mechanisms that underlie molecular aging. We therefore quantified the concerted activity of the full aging signature at the single cell level. To assess the biological utility of the intrinsic aging signature in explaining cellular heterogeneity in a neurodegenerative context, we computed the systemic and intrinsic signature activity score per cell in a dataset comprising AD patients and age-matched controls (age range 49-92) ^38^. Neurons with high/low systemic age signature score were homogeneously distributed and did not exhibit cell-type-specific enrichment. Nevertheless, neurons from AD patients were predicted to be older than those from healthy controls based on systemic age signature (Figure 3E, Top), consistent with the accelerated brain aging observed in AD ^39^. In contrast, intrinsic age signature revealed pronounced cell type heterogeneity, which was further amplified in AD. Inhibitory neuronal populations were enriched for cells with low intrinsic age signature score in healthy controls, but exhibited a marked loss of these molecularly younger cells in AD relative to controls (Figure 3E, Bottom).

Extending cortical aging signatures to other brain regions and disease states, we examined SNc dopaminergic neurons in LBD and PD patients ^40^. We calculated the activity scores in dopaminergic neurons from the substantia nigra pars compacta (SNpc) in Lewy body dementia (LBD) and Parkinson’s disease (PD) patients ^40^. In LBD, similar to AD, neurons displayed accelerated aging compared to controls. In PD, profiling aging in dopaminergic neurons revealed distinct patterns for cell types (Extended Data Figure 7C). Notably, cell types that appeared molecularly old based on systemic age in controls –particularly the SOX6-DDT subtype– were selectively depleted in PD. At baseline, SOX6-DDT exhibits high expression of intrinsic age-associated oxidative phosphorylation genes MT-CO1 and COX5B, while showing low expression of a dopaminergic vulnerability marker, Regulator of G protein signaling 6 (RGS6) which has been suggested to be neuroprotective ^41^. In contrast, SOX6-AGTR1, a subtype previously described as vulnerable, displayed the opposite aging patterns, with high RGS6 expression and low expression of aging related genes at baseline. The depletion of both SOX6-AGTR1 and SOX6-DDT in PD supports the notion that intrinsic, age-associated molecular features at baseline reflect cell type-specific mechanisms that influence survivability upon exposure to pathology. Together, these results highlight the heterogeneity of cellular aging responses across distinct disease contexts, underscoring that transcriptional aging signatures capture biologically meaningful differences in vulnerability across cell types rather than a uniform aging trajectory.

### Intrinsic Aging is Predictive of Cellular Vulnerability and Resilience

Next, we investigated the utility of the intrinsic aging signature in predicting cellular vulnerability at the cell-type level. The ratio of each cell type was calculated as its proportion within a subject group relative to its proportion in the total cell population. Cell types with a ratio below 1 were classified as underrepresented (vulnerable), while those with a ratio above 1 were considered overrepresented (resilient) in the group of interest, with the reference group serving as the baseline. In-SST interneurons exhibited the lowest ratio in the old subject group, while Ex-L23 exhibited the highest ratio denoting an increase in its proportion as a result of the decline in other cell populations with age (Figure 4A).

**Figure 4.**
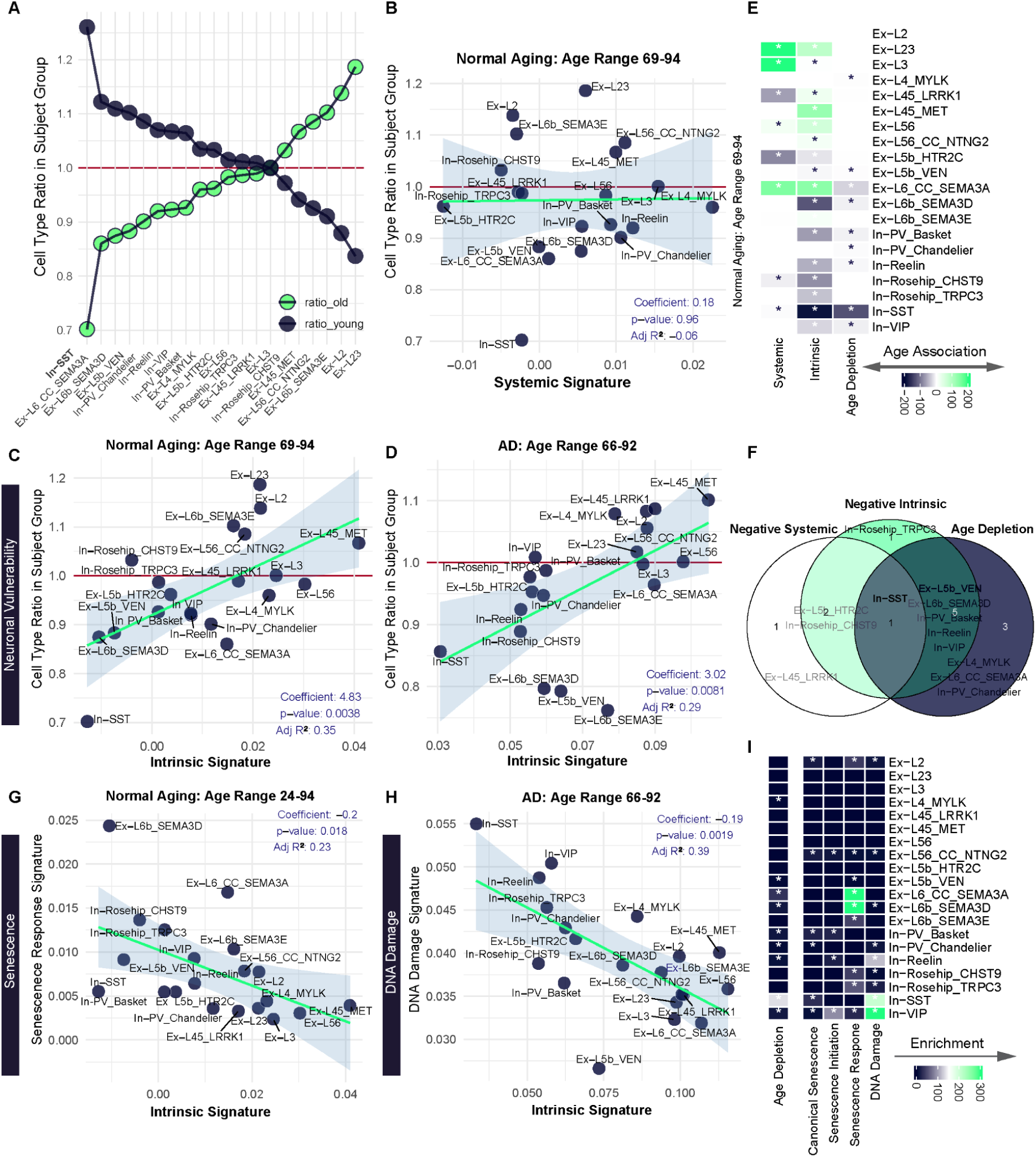
Cell-Type differential vulnerability in healthy aging and Alzheimer’s disease. A. Ratio of each cell type relative to other cell types in young and old subject groups, highlighting In-SST to be the most depleted in older subjects. B,C. Relation of the mean systemic (B) and intrinsic (C) aging signature activity score per cell type (x-axis) to the ratio of a cell type in old subjects with age range 69-94 years (y-axis). D. Relation of the mean intrinsic aging signature activity score per cell type (x-axis) to the ratio of a cell type in AD patients with age range 66-92 years (y-axis). E. Heatmap for the overrepresentation of cells enriched for aging gene signatures per cell type and underrepresentation (depletion) of a cell type in old subjects. Color scale indicates –log₁₀(*p*) for the hypergeometric test multiplied by +1 for gene sets positively associated with systemic or intrinsic age and –1 for gene sets negatively associated with age. For cell types underrepresented with age, the –log₁₀(*p*) is multiplied by –1 to reflect depletion. *P*-values were adjusted for multiple comparisons using false discovery rate (FDR) correction applied across all cell types (rows) and features/conditions (columns); adjusted *p*-values are displayed on the plots. F. Venn Diagram for cell types enriched for signatures negatively associated with systemic or intrinsic age and depleted with age based on hypergeometric test, highlighting In-SST to be depleted with age and enriched for negative systemic and intrinsic signatures. G. Relation of the mean intrinsic aging signature activity score (x-axis) and the mean senescence response signature activity score (y-axis) and per cell type in the full cohort of healthy subjects with age range 24-94. H. Relation of the mean intrinsic aging signature activity score (x-axis) and the mean DNA damage signature activity score (y-axis) per cell type in AD subjects, highlighting the highest DNA damage score for In-SST more than expected based on its cellular age. I. Heatmap for the overrepresentation of cells enriched for positively or negatively associated aging signatures, canonical senescence, senescence initiation, senescence response and DNA damage signatures per cell type and underrepresentation of a cell type in old subjects. Color scale indicates –log₁₀(*p*) for the hypergeometric test. *P*-values were adjusted for multiple comparisons using false discovery rate (FDR) correction applied across all cell types (rows) and features/conditions (columns); adjusted *p*-values are displayed on the plots. In B-D and G-H. Each point represents a cell type. Linear regression was used to assess associations; regression coefficients, *p*-values, and adjusted R² are displayed on the plots.

Paradoxically, although we expected molecularly older neurons (higher intrinsic age) to exhibit higher degenerative rates (lower ratio in older subjects), we found the opposite to be true. By regressing the cell type ratio on the mean aging signature score, we identified a significant positive association between the ratio and the mean intrinsic age signature score in older subjects (69–94 years). However, this association was not significant for the systemic age signature (Figure 4B-C). Overall, inhibitory cell types exhibited a lower ratio and a lower mean intrinsic age compared to excitatory cell types, particularly upper layer excitatory neurons. As expected, In-SST exhibited the lowest mean intrinsic age with its lowest ratio.

To confirm the observed cellular vulnerability, we applied a hypergeometric test to each cell type. Specifically, two tests were conducted per cell type to assess the overrepresentation of cells enriched for genes positively or negatively associated with intrinsic age. If a cell type was more enriched for the positively associated features, the p-value from the overrepresentation test was assigned a positive sign; if it was more enriched for the negatively associated features, the p-value was assigned a negative sign. For cell type composition, underrepresentation was interpreted as depletion with age or disease, and accordingly, the significance level was also assigned a negative sign. As expected, positive intrinsic age-associated signature was more enriched in cells isolated from old relative to young individuals (Figure 4E) and the opposite trend was observed for intrinsic negatively age-associated signature. Similar to the ratio test, inhibitory neurons and certain deep-layer excitatory neurons were underrepresented in the old group, indicating their age-related vulnerability (Figure 4E, 3^rd^ column). This was reflected in their negative association with the intrinsic age signature, which denotes an overrepresentation of neurons enriched for genes negatively associated with intrinsic age (Figure 4E, 2^nd^ column), but not for the systemic age signature. Five cell types depleted with age showed exclusive enrichment for the negative intrinsic signature, whereas only one –In-SST– was depleted with age and enriched for the negative systemic and intrinsic signature (Figure 4F).

To further explore the utility of this approach in understanding neuronal vulnerability to neurodegeneration, we applied it to the AD dataset ^38^. Similar to aging, a positive association between the ratio and the mean intrinsic age signature score in AD patients was observed (Figure 4D). In-SST still exhibited low ratio in AD but was preceded by deep layer excitatory neurons: Ex-L6b-SEMA3E, Ex-L5b-VEN and Ex-L6b-SEMA3D, which exhibited lower ratio than expected based on their mean intrinsic age (Extended Data Figure 5F). Additionally, we observed a similar pattern of depletion to aging: inhibitory neurons were underrepresented in AD patients and negatively associated with the intrinsic age signature (Extended Data Figure 5H, 2^nd^-3^rd^ columns). Four AD-depleted cell types were exclusively enriched for the negative intrinsic signature, including In-SST. The only AD-depleted cell type enriched for both negative systemic and intrinsic signatures was Ex-L5b-VEN (Extended Data Figure 5I).

Among inhibitory neurons, we also observed associations between neuron vulnerability and intrinsic age. In healthy aging, In-PV (parvalbumin) and In-VIP (vasoactive intestinal polypeptide) were underrepresented in the aged individuals compared to the young, however, this pattern was not observed in AD. Although this may appear contradictory, it reflects differences in the statistical context in which these comparisons were made. When considering total neuronal population across age-matched controls and AD patients, these cell types do not appear selectively depleted in AD because their reduction has already occurred during normal aging. In contrast, In-Reelin and In-SST were underrepresented in both aging and AD, consistent with prior reports ^42,43^ and supporting the heightened vulnerability of these cell types in AD. Thus, neurons with low/negative intrinsic age (molecularly younger) are the most vulnerable with heightened OXPHOS activity. These findings confirm our initial observations that high/positive intrinsic age is associated with neuronal resilience to aging and neurodegeneration while low/negative intrinsic age is associated with vulnerability.

### Intrinsic Aging is Predictive of the Response to Senescence and DNA Damage

Based on the observation that the enrichment for the intrinsic aging signature marks cellular vulnerability, we hypothesized that this signature might also reflect key hallmarks of aging, such as senescence and DNA damage. Leveraging previously published gene signatures for canonical senescence, senescence initiation and senescence response ^44^, we observed a significant inverse association between mean senescence response score and mean intrinsic age signature score across healthy subjects spanning 24-94 years of age, again, this was not observed with mean systemic age signature (Figure 4G, Extended Data Figure 6A-B). At the cell-type level, most excitatory neurons exhibited higher senescence response relative to their intrinsic age score, with Ex-L6b-SEMA3D displaying the strongest senescence response. In contrast, inhibitory neurons exhibited lower senescence response relative to their intrinsic age score. Given the positive association of the intrinsic age signature score with cell type ratios, together with their negative association with senescence response signature score, we hypothesized that senescence response would be inversely associated with cell-type abundance irrespective of disease status. Contrary to this expectation, we observed a disease-specific pattern: senescence response negatively associated with the cell type ratios only in AD, yet positively associated in age-matched controls (Extended Data Figure 6C-D). Consistent with this divergence, Ex-L6b-SEMA3D exhibited the highest abundance in controls and lowest abundance in AD patients.

Similarly for DNA damage ^45^, mean DNA damage signature score was inversely associated with the mean intrinsic age signature, but not with the mean systemic age signature score. This relationship was observed for both aged healthy subjects (Extended Data Figure 6E-F) and AD patients (Figure 4H, Extended Data Figure 6G-H), but was more pronounced in the AD state. In-SST had the highest DNA damage response with the lowest mean intrinsic age, followed by In-Reelin and In-VIP.

To integrate cellular heterogeneity incurred by the response to senescence and DNA damage with intrinsic aging, we quantified deviations in the proportion of cells enriched for these signatures across subject groups and cell types. Neurons enriched in DNA damage and senescence initiation signatures were overrepresented in old subjects relative to young subjects (Extended Data Figure 6I) and in AD patients relative to age-matched controls (Extended Data Figure 6J). Meanwhile, canonical senescence was selectively enriched in healthy aged controls, whereas senescence response was not enriched in any group (Extended Data Figure 6I-J). This underscores the indispensability of the cell-type-resolved analyses for understanding aging hallmarks, which become evident only upon stratifying enriched cells into the cell types within healthy old and AD groups. Consistent with earlier analyses, senescence response signature was most significantly enriched in deep layer excitatory neurons Ex-L6-CC-SEMA3A and Ex-L6b-SEMA3D in healthy aging and AD (Figure 4I, Extended Data Figure 6K-L), also shown in (Extended Data Figure 6A-B). In contrast, DNA damage signatures were most prominently enriched in inhibitory neuron subtypes (Figure 4I, Extended Data Figure 6K-L), also shown in (Extended Data Figure 6E-F). These results further reinforce the hypothesis that lower intrinsic aging is associated with neuron vulnerability, as well as, other hallmarks of aging depending on the cell type identity.

### Intrinsic Aging is Predictive of Epigenetic Erosion

Next, we sought to evaluate changes of the regulatory landscape in the form of epigenetic erosion: loss of epigenetic demarcations in cellular aging. We quantified epigenetic erosion score per cell in the AD dataset ^38^ as previously described ^46^, where cells with positive score exhibit higher chromatin accessibility in normally repressed regions and cells with negative score exhibit higher chromatin accessibility in normally open regions.

Across cell types, excitatory neurons exhibited more positive epigenetic erosion scores relative to inhibitory neurons (Figure 5A). This was evident at the neuron subtype level as well, where inhibitory subtypes displayed more negative epigenetic erosion scores. Across conditions and consistent with prior reports, we confirmed a global increase in epigenetic erosion in AD patients relative to healthy controls ^46^. To determine whether this effect was cell-type-specific, we next compared erosion score distributions between AD and controls across cell types. Interestingly, we were able to attribute this effect primarily to inhibitory neurons. Inhibitory neurons exhibited a significant shift toward higher accessibility of repressed regions in AD, whereas excitatory neurons exhibited a shift toward reduced accessibility (Figure 5B, two-sample Kolmogorov-Smirnov test (KS), D=0.15 for inhibitory neurons and D=0.07 for excitatory neurons). Upon stratifying neurons in subtypes, we observed more pronounced shifts in erosion. In-Reelin and In-PV-Basket displayed marked positive shifts in accessibility in AD relative to controls (Extended Data Figure 8D, KS D=0.3 and KS D=0.17, respectively). Ex-L6b-SEMA3D, a highly senescent and AD-depleted cell type, exhibited a more concentrated erosion score distribution toward more positive scores in AD (Extended Data Figure 8D, KS D=0.13), counter to the broader excitatory trend.

**Figure 5.**
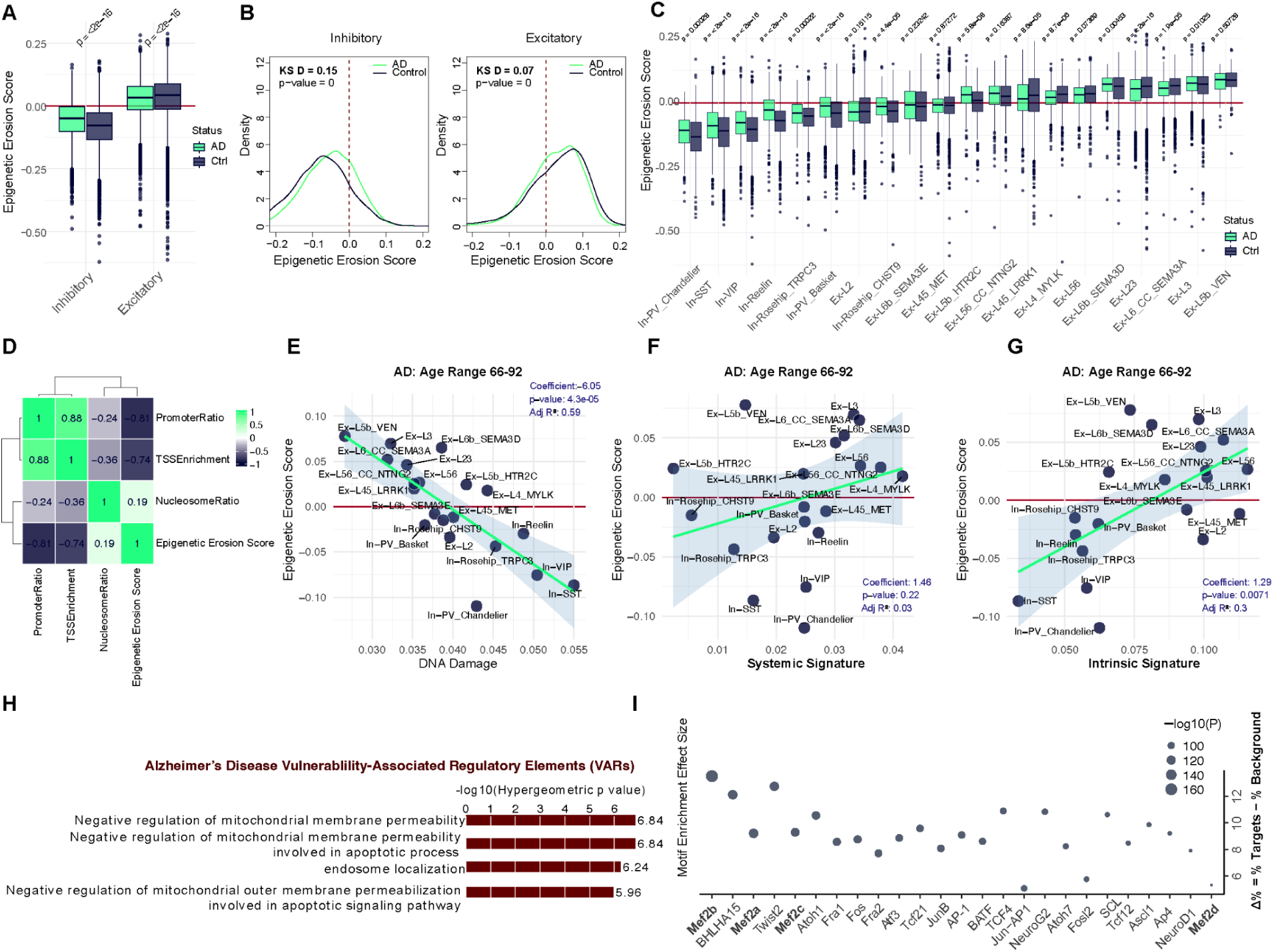
Distributional and cell-type-specific differences in epigenetic erosion scores in healthy aging and Alzheimer’s disease. A. Boxplots for epigenetic erosion score across excitatory and inhibitory neurons in AD patients relative to age-matched controls. Distributions are shown separately for each cell type and subject group. Center lines indicate medians, boxes represent interquartile ranges, and whiskers extend to 1.5×IQR. Statistical comparisons between AD and control groups were performed independently within each cell type using two-sided Wilcoxon rank-sum tests (Mann-Whitney U tests; unpaired, non-parametric). Each observation represents a single cell. Exact raw *p*-values are displayed on the plots. B. Density distributions of epigenetic erosion score for excitatory (left) and inhibitory (right) neurons in AD patients relative to age-matched controls. Curves represent kernel density estimates of single-cell epigenetic erosion scores for each group. Distributional differences between AD and control cells were assessed within each cell type using two-sample Kolmogorov-Smirnov tests (two-sided, non-parametric). Each observation represents an individual cell. KS statistics (D) and exact raw p-values are shown on the plots. Red dashed line marks zero. C. Boxplots for epigenetic erosion score across neuronal cell subtypes in AD patients relative to age-matched controls. Distributions are shown separately for each cell subtype and subject group. Center lines indicate medians, boxes represent interquartile ranges, and whiskers extend to 1.5×IQR. Statistical comparisons between AD and control groups were performed independently within each cell type using two-sided Wilcoxon rank-sum tests (Mann-Whitney U tests, unpaired, non-parametric). Each observation represents a single cell. Exact raw *p*-values are displayed on the plots. D. Hierarchically clustered pairwise Pearson’s correlations between epigenetic erosion score, nucleosome ratio, promoter ratio and TSS enrichment. Correlations were computed across single cells. E-G. Relation of the DNA damage signature activity score (E), mean systemic aging signature activity score (F) (x-axis) or mean intrinsic aging signature activity score (G) and the mean epigenetic erosion score (y-axis) per cell type in AD. Each point represents a cell type. Linear regression was used to assess associations; regression coefficients, *p*-values, and adjusted R² are displayed on the plots. H. Gene ontology enrichment analysis of AD vulnerability-associated regulatory elements (VARs), which are the differentially accessible chromatin regions in inhibitory neurons enriched for the intrinsic aging signature that is negatively associated with age. Differential chromatin accessibility was assessed using limma-voom, regions in the 98% quantile of the logFC were filtered, followed by adjusting p-values and weighted FDR < 0.05 threshold was set. Enriched gene ontology terms were identified using GREAT. Foreground set: 3,362 distal differentially accessible chromatin regions, Background set: 172,210 distal accessible chromatin regions across cell types and conditions. I. Transcription factor motif enrichment analysis of open chromatin regions defined in panel H. The top 25 enriched transcription factor motifs with *p* < 1 × 10⁻³⁰ are shown. Motif enrichment effect size is reported as the difference in the percentage of target regions versus background regions containing each motif (Δ% = % targets − % background; y-axis). Statistical significance is displayed as −log₁₀(*p*) and reflected by point size.

Given the pronounced erosion shifts and vulnerability of inhibitory neurons during both healthy aging and AD, we focused on this population. We binned inhibitory cells depending on gene set signature scores. In AD, the candidate vulnerable neurons enriched for lower intrinsic age scores and high DNA damage signatures exhibited higher promoter ratio and TSS enrichment, despite lower number of fragments (Extended Data Figure 8B-C). No comparable differences were captured by the systemic age signature. We then related these signatures to erosion at the cell-subtype level. Mean DNA damage signature score was negatively associated with mean erosion (Figure 5E), consistent with prior reports of downregulated DNA damage and repair processes in neurons with high epigenetic erosion ^46^. While, intrinsic aging was positively associated with epigenetic erosion, underscoring the granularity of the signature in predicting chromatin state based on cellular aging profile. These associations place the vulnerable inhibitory population –marked by low intrinsic age and high DNA damage– at low baseline erosion, yet it is precisely these cells whose distribution shifts toward more positive scores in AD.

### Machine Learning Models Reveal Intrinsic Age as the Most Informative Feature for Cellular Vulnerability

To evaluate the relative contribution of distinct molecular signatures to cellular vulnerability, we classified cells into vulnerable or resilient cells based on whether the cell type is underrepresented in the subject group of interest. Activity signature scores for seven quantified phenotypes: systemic age, intrinsic age, DNA damage, canonical senescence, senescence initiation, senescence response and epigenetic erosion, were utilized as features to train three binary classifiers to predict whether a neuron is vulnerable or resilient. Models were trained on a designated training set and evaluated on an independent test set, with all performance metrics computed exclusively on the test set. Decision tree complexity was optimized using the 1-SE rule based on cross-validated error.

In the context of normal aging, intrinsic age was positioned at the root node of the decision tree, with epigenetic erosion and senescence response in the downstream branch nodes (Extended Data Figure 9A). We additionally trained Random Forest and XGBoost to benchmark their performance against the decision tree. All three classifiers demonstrated comparable performance, with Random Forest and XGBoost achieving higher AUC values for the ROC curve, at 0.85 (Extended Data Figure 9B). A similar pattern was observed in AD, where intrinsic age again emerged at the root node (Extended Data Figure 9D). Random Forest and XGBoost continued to outperform the decision tree with AUCs reaching 0.88 and 0.87, respectively (Extended Data Figure 9F).

Notably, across both aging and AD contexts, intrinsic age consistently emerged as the most informative feature, as evidenced by its placement at the root node of the decision tree, the largest mean decrease in accuracy when excluded from the Random Forest model (Extended Data Figure 9B,E) and the highest gain in the XGBoost model.

### Regulatory Elements Linking Mammalian Longevity and Human Age Vulnerability

To identify the open chromatin regions associated with vulnerability in inhibitory neurons, we examined chromatin accessibility in cells negatively associated with the intrinsic aging signature in AD patients. Differentially accessible regions in vulnerable neurons were termed as **vulnerability-associated regions (VARs)**. Using a 98th-percentile logFC threshold, we identified 3,362 differentially open peaks or VARs (positive logFC) and 3,362 differentially closed peaks (negative logFC) in vulnerable neurons (Supplementary Table 5). Open peaks were defined by the 98th-percentile cutoff (logFC ≥ 1.4, adjusted p ≤ 1e-12) and closed peaks by the 2nd-percentile cutoff (logFC ≤ −1.4, adjusted p ≤ 2e-12). Gene ontology analysis using GREAT revealed that VARs were significantly enriched for pathways related to negative regulation of mitochondrial membrane permeability in apoptotic signaling and endosome localization (Figure 5H). This aligns with earlier observations that mitochondrial functional status is a key determinant of neuronal vulnerability, as observed in genes negatively associated with intrinsic age (Figure 3B). In contrast, accessible regions in neurons enriched for systemic aging failed to capture the same intrinsic processes. Transcription motif enrichment analysis of VARs highlighted the Mef2 family of transcription factors (Mef2a, Mef2b, Mef2c and Mef2d, p < 10e-30, Figure 5I), critical regulators of neuronal resilience to neurodegeneration ^47,48^.

To investigate the relationship between longevity and aging, we asked whether there is a consensus on critical regulatory elements across mammals and within the human population. For aging vulnerability, we used a more permissive 90th-percentile threshold, we identified 16,809 open (VARs) and 16,809 closed peaks, with open peaks defined by the 90th-percentile cutoff (logFC ≥ 1, adjusted p ≤ 3e-05) and closed peaks by the 10th-percentile cutoff (logFC ≤ −0.8, adjusted p ≤ 1e-05). For longevity, we pooled LARs across cortical cell types –including neurons, glia, and endothelial cells– based on direction of association, yielding non-overlapping 8,447 positively and 12,134 negatively associated unique peaks. Notably, the differentially open peaks that distinguish the vulnerable neurons significantly overlap the negative LARs, which have lower predicted open chromatin in longer-lived species (adjusted p-value ≤ 0.05). The 427 unique shared regions that make up this intersection are termed as **vulnerability and longevity regions (VLRs)** (Figure 6A-B, Supplementary Table 6). At the cell-type level, this enrichment was most pronounced in In-PV neurons, indicating that this population contributes substantially to the global signal, followed by In-SST and Ex-L5-ET neurons. While positive LARs also showed overlap (Figure 6B, left), negative LARs exhibited both greater overlap and stronger enrichment with the OCRs that mark vulnerable neurons (Figure 6B, right). This asymmetry suggests that there is selective pressure to down-regulate open chromatin associated with neuron vulnerability in species with longer lifespans.

**Figure 6.**
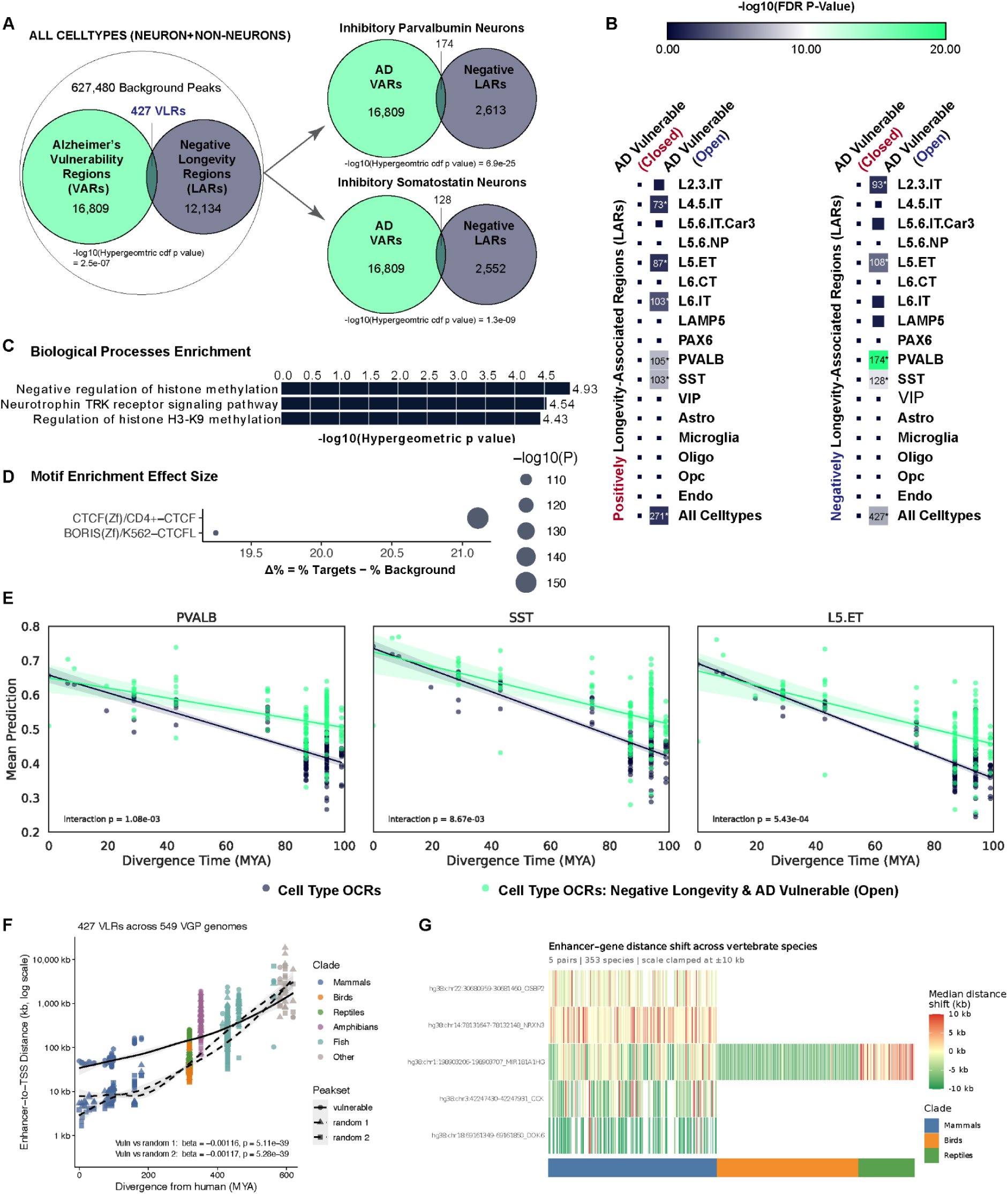
Shared Molecular Pathways of Vulnerability in Mammalian Longevity and Human Aging and Conservation Across Vertebrates. A. Common regulatory elements between vulnerable neurons open chromatin regions (VARs) and negatively longevity-associated regions (LARs), resulting in 427 vulnerability and longevity regions (VLRs). B. Enrichment of overlap between AD vulnerable neurons open/closed chromatin regions and positively/negatively longevity-associated regions (LARs) across 17 cortical cell types. Color scale and square size represent enrichment significance (−log10 FDR-adjusted P-value) based on the complement of the hypergeometric cumulative distribution function. Box sizes are capped to emphasize significant enrichments. C. Gene ontology enrichment analysis of the 427 VLRs shared regulatory elements (foreground), with the combined set of VARs and negative LARs used as background (GREAT). D. Transcription factor motif enrichment analysis of 427 VLRs shared regulatory elements. E. Mean predicted enhancer activity as a function of divergence time from humans (millions of years ago, MYA) across 240 mammalian genomes from the Zoonomia database. Each point represents the mean predicted activity across all enhancers in the corresponding set for a given species. Solid lines indicate linear regression fits with 95% confidence intervals. Vulnerability-associated regions, defined as enhancers overlapping cell-type-specific negatively longevity-associated and negatively aging-associated peaks in PVALB interneurons, SST interneurons, and L5-ET neurons (mint green), are compared with non-vulnerable background cell-type-specific regulatory regions (navy). Interaction *P*-values correspond to the divergence time × enhancer class interaction term from ordinary least squares regression models (***mean predicted activity ∼ divergence time × enhancer class***), testing whether the relationship between divergence time and predicted enhancer activity differs between enhancer sets. F. Median enhancer-gene distance relative to the human reference genome (hg38) as a function of divergence time from humans (millions of years ago, MYA) across 577 vertebrate genomes from the Vertebrate Genomes Project (VGP). Each point represents a species and is colored by major vertebrate clades. The 427 VLRs (solid triangles, dashed trend line) are compared with two size-matched sets of non-overlapping randomly sampled longevity negative regions (solid circles, solid trend line). Black curves indicate LOESS fits with 95% confidence intervals. Interaction *P*-values correspond to the divergence time × enhancer class interaction term from linear regression models (***log10(median distance + 1) ∼ divergence time × enhancer class***), testing whether the relationship between evolutionary divergence and enhancer-gene distance differs between vulnerable and background enhancer sets. G. Median enhancer–gene distance relative to the human reference genome (hg38) for 7 syntenic VLR-gene pairs across 353 species and 3 clades.

Functional enrichment analysis of the shared VLRs revealed pathways involved in negative regulation of histone methylation (Figure 6C). This is consistent with the dynamic nature of histone methylation, which is regulated by the opposing activities of methyltransferases and demethylases. These enzymes have been reported as critical and evolutionarily conserved regulators of longevity, and their dysregulation has been shown to alter tissue aging and lifespan across species ^49^. Neurotrophin Trk receptor signaling was also enriched in the shared regulatory elements. Neurotrophin pathways are key regulators of neuronal survival and synaptic plasticity. Particularly, downregulation of the BDNF-TrkB signaling axis has been implicated in aging and age-related cognitive disorders ^50^. Motif analysis further revealed enrichment of the CTCF motif (Figure 6D, p = 10e-66), as well as MEF2B (p = 1e-13). CCCTC-binding factor (CTCF) is a highly conserved architectural protein that organizes 3D genome structure ^51^.

Consistent with their potential functional importance, VLR peaks were more evolutionarily conserved across mammals than background cell-type-specific open chromatin regions. This difference was supported by significant interaction between divergence time and enhancer group in a linear regression model for PVALB, SST, and L5-ET neurons independently (Figure 6E).

To determine whether this pattern extended beyond mammals, we leveraged the 577-species from the Vertebrate Genome Project (VGP) ^52^ alignment. We first assessed nucleotide-level conservation by computing average phyloP scores per region^52,53^. Conservation was significantly higher in vulnerable open regions than in closed regions, and in negative-longevity regions than in positive ones, though both shifts were subtle (Extended Data Figure 10A-B). We next compared VLRs against two equal-sized sets of randomly sampled regions drawn from negative LARs (15,151 regions). VLRs showed no significant increase in conservation relative to either set (Extended Data Figure 10C). This was consistent with our expectation, since these regions were prioritized based on learned enhancer activity conservation rather than nucleotide-level sequence conservation ^10^. One way to evaluate conservation of enhancer function is through its proximity to a specific gene. We therefore examined conservation of enhancer-gene syntenic relationships across vertebrates. For each enhancer, we quantified the distance to its nearest gene and evaluated how this relationship changed with increasing evolutionary divergence from humans. As expected, enhancer-gene distances generally increased with divergence time, consistent with the accumulation of genomic rearrangements over evolutionary timescales. However, VLRs exhibited significantly slower divergence in enhancer-gene distance than two non-overlapping matched sets of randomly sampled negatively LARs (Figure 6F). While VLRs were initially located farther from their target genes in closely related species, their enhancer-gene distances increased more gradually across vertebrate evolution, indicating greater conservation of syntenic relationships. Again, the interaction between divergence time and enhancer group confirmed the conservation (beta = −0.00116, p = 5.11e-39 for random set 1 and beta = −0.00117, p = 5.28e-39 for random set 2).

We next focused on VLRs that maintained syntenic relationships with vulnerability-associated genes across species (Figure 6G). This analysis identified VLRs near to *MIR181A1HG, CCK, NRXN3, DOK6, and OSBP2*. Notably, *MIR181A1HG* was the only locus for which enhancer-gene synteny was preserved across mammals, birds, and reptiles, with an orthologous enhancer retained on chromosome 1 throughout these lineages. In contrast, syntenic enhancer-gene relationships involving *CCK, NRXN3, DOK6, and OSBP2* were restricted to mammals. The co-occurrence of a synaptic adhesion gene (*NRXN3*)^54^, a major neuromodulator (*CCK*)^55^, a mediator of neurite growth (*DOK6*)^56^ and a posttranscriptional regulator of synaptic plasticity (*MIR181A1HG*)^57^ points to molecular pathways supporting synaptic plasticity, connectivity, and circuit maintenance. The negative association of these genes with aging, together with their proximity to VLRs, suggests that evolutionarily conserved regulatory elements may contribute to age-related declines in neuronal maintenance and synaptic function.

## DISCUSSION

The age dependence and cell-type-specificity of NDs hold promise for the development of therapeutics that selectively target vulnerable cell populations. However, therapeutic development is hindered by the complex and heterogeneous etiologies of NDs, which often lack well-defined causal drivers ^58^. To address this, we leveraged comparative genomics across placental mammals and brain single-cell data to prioritize the specific genes and regulatory elements underlying selective vulnerability to aging and NDs.

In this study, we built a multi-scale framework to deconvolve cell-type-specific aging and vulnerability. At the level of placental mammals, we resolved longevity-associated selective pressure by applying the TACIT method to the Zoonomia database, and extended our analysis across vertebrates from the VGP database. At the level of the human population, we developed a transcriptional aging clock that yielded a cell-type-specific vulnerability metric to flag vulnerable neurons and identify the regulatory networks associated with such selective vulnerability. We find that intrinsic cellular age, driven by stress-response factors and dysregulation of mitochondrial oxidative phosphorylation, is a stronger predictor of cell loss during both normal aging and Alzheimer’s disease progression and tracking more closely with DNA damage, senescence and epigenetic erosion. Spanning different methods and scales, these independent analyses converged on an epigenomic signature of neuronal vulnerability under differential selective pressure across mammals (Figure 1).

Longevity varies greatly across placental mammals, providing an opportunity to find loci that are candidate drivers of aging. Previous approaches have noted longevity-associated differences in genes^59^ and DNA methylation^16,60^. We extend this work to longevity-associated differences in regulatory elements. Directly measuring OCRs is intractable in a consistent tissue across hundreds of species. However, we are able to overcome this limitation by using CNNs models trained in four species (human, macaque, mouse, and rat) to predict chromatin accessibility (activity) across 240 placental mammals. We find 11,379 positively- and 15,151 negatively-associated open chromatin regions. These OCRs may function as enhancers, insulators, or other types of *cis*-regulatory elements that control gene expression levels. LARs were enriched to be near gene ontology categories that have been linked to aging and longevity evolution, including histone modification, cellular stress, and mitochondrial function. Both within and across cell types, multiple pathways at the intersection of cell intrinsic apoptosis and DNA damage emerged as being enriched, an intersection directly tied to aging^61^. This result demonstrates that there are selective pressures in longer-lived species to reduce or increase open chromatin levels near genes associated with longevity pathways.

Traditional methods for associating genetic differences across species with complex traits, like aging, rely on finding signals in individual nucleotides ^62^. In contrast, the cell-TACIT method has the ability to stratify the evolutionary pressures across specific cell types based on the activity of the regulatory element. This allowed us to test the hypothesis that different hallmarks of aging are under selective pressure in different types of cortical cells. Indeed, we found that the OCRs inferred across different cell types were enriched to be near genes related to different hallmarks of aging, often related to the function of the cell type. For example, inferred LARs for microglia, brain resident immune cells, were near genes related to inflammation and the response to misfolded proteins. In contrast LARs in oligodendrocytes were near genes related to myelination, which has also been associated with species longevity. Rather than a few pathways that are master regulators of longevity, our results suggest a model in which different pathways are under selective pressure for longevity depending on the constraints that longer life spans put on that particular cell type (Figure 2).

We then set out to test whether the longevity-associated OCRs could relate directly to selective neuron vulnerability to aging and ND in the human brain. We investigated age-associated gene expression signatures across the human population using published cohorts of single-cell gene expression and open chromatin from post-mortem brain tissue. As our longevity analysis implicated cell intrinsic mechanisms, we applied a machine learning framework to those datasets to distinguish cell intrinsic mechanisms of aging versus those that are systemic across the brain tissue of healthy individuals. Relative to systemic aging signatures, we find that cellular intrinsic age is a stronger predictor of cell loss during aging and the progression of AD. Moreover, intrinsic age is more associated with other hallmarks of aging, including DNA damage, senescence and epigenetic erosion. The model was also able to capture phenomena such as accelerated brain aging in AD patients ^39^, with AD-derived neurons predicted to be systemically older than those from healthy controls and did not exhibit cell-type-specific patterns. In contrast, the intrinsic aging model was able to resolve subtype-specific differences across normal aging and disease states.

At the outset of the study, we expected to find intrinsic cellular age to be positively associated with key pathological features underlying aging and NDs. Paradoxically, we found that molecularly younger neurons are the most vulnerable, low-intrinsic-age neurons undergo selective loss with heightened responsiveness to senescence, DNA damage or epigenetic erosion, in addition to disrupted mitochondrial bioenergetics. Vulnerable neurons exhibited elevated expression of nuclear-encoded OXPHOS programs, further reinforcing mitochondrial dysfunction, a central feature of aging and pathophysiology of neurodegenerative diseases ^63,64^. This imbalance was also evident at the regulatory level: in AD, regulatory networks governing mitochondrial membrane permeability were selectively active in these vulnerable cell types, however, this enrichment was not observed in controls. This regulatory landscape was further characterized by enrichment of MEF2 transcription factor motifs, which are key regulators of neuronal resilience to neurodegeneration ^47^. This suggests that neuron subtypes with higher intrinsic age are, in fact, the more adaptive and resilient, upregulating protective gene programs as they age, whereas subtypes with lower intrinsic age express programs that are downregulated with age, eroding their functional identity and accelerating their vulnerability to aging. Together, these results support a model in which aging and neurodegenerative diseases impose bioenergetic challenges on neuron subtypes across the brain: subtypes able to meet those challenges by inducing neuroprotective programs are more likely to survive, while those unable to respond are more likely to be lost. Therapeutics that recalibrate these expression programs in vulnerable neuron subtypes might thereby slow the progression of neurodegeneration.

To evaluate the evolutionary conservation of the identified regulatory programs underlying cellular vulnerability in human aging, we leveraged the cell-type-aware cross-species framework, enabling the identification of shared molecular mechanisms linking mammalian lifespan variation to human neuronal aging. Notably, differentially active regions in vulnerable neurons overlapped strongly with LARs in mammals, particularly in the negative direction, highlighting common regulatory mechanisms underlying neuronal vulnerability in humans and shorter lifespans across mammals.

The convergence of negatively longevity-associated and age-vulnerability regulatory elements (VLRs) within vulnerable neuronal populations, In-PV, In-SST and L5-ET offer a molecular bridge between evolutionary constraints on lifespan and human brain aging. Interestingly, a circuit model has been proposed where layer 5 ET forms reciprocal connectivity with inhibitory PV and SST^65^, and here we extend that this circuit motif is under selective pressure. This further validates the ability of the intrinsic aging clock to selectively identify neuronal populations most susceptible to aging. The enrichment of VLRs near genes implicated in synaptic adhesion, neuromodulation, and plasticity suggests that regulatory deterioration at these loci may represent a conserved mechanism through which aging erodes circuit integrity. Conservation of syntenic enhancer-gene relationships across vertebrates, particularly at the *MIR181A1HG* locus, together with CTCF enrichment in VLRs, confirms that chromatin organization and regulatory architectures are under sustained selective pressure.

Our findings suggest that the chromatin accessibility landscape of vulnerable neurons is itself a target of evolutionary constraint linked to longevity. The open chromatin regions most strongly associated with neuronal vulnerability tend to be under negative selection, and their accessibility is negatively correlated with species lifespan. In other words, the regulatory elements that are accessible in vulnerable neuronal populations appear to be subject to evolutionary pressure to be closed or attenuated, in longer-lived species. We therefore propose a causal link extending from cell-type-specific regulatory failure to cellular vulnerability and, ultimately, to shorter lifespans within and across species.

Future work may investigate whether leveraging larger single-nucleus human brain cohorts further improves iENR model performance or enables the discovery of additional biological insights. In this present study, we intentionally trained our elastic net regression model using a moderately-sized cohort of 69 individuals ^20^, selected to achieve a balance of broad lifespan coverage, technical homogeneity, statistical power and computational efficiency. Another important consideration, elastic-net models perform optimally in high-dimensional, low-sample settings (p >> n) and are sensitive to confounders, including technical heterogeneity and batch effects that can be amplified when aggregating heterogeneous datasets to increase cohort size. Given the wide adult age span (24-94 years), highly resolved neuronal cell type profiles, the selected dataset provided sufficient statistical power to learn robust neuron-specific age associated signals, while preserving interpretability and biological specificity.

Altogether, we built a multi-scale framework that spans evolutionary-extrinsic, tissue-systemic and cellular-intrinsic aging effects to profile the candidate causal drivers of cellular vulnerability. This work introduces a generalizable and interpretable framework for decomposing systemic and cell-specific aging signals in single-cell data, enabling resolution of cellular heterogeneity across disease contexts. It demonstrates that transcriptional aging signatures encode biologically meaningful differences in vulnerability. By linking intrinsic aging to cellular vulnerability and evolutionary conservation, it provides a foundation for prioritizing therapeutic targets and understanding mechanisms underlying selective neurodegeneration, offering a means to preserve vulnerable cell types and limit or slow the progression of neurodegenerative diseases. More broadly, it offers a scalable approach for systematic interrogation of diverse phenotypes at the single-cell resolution and is readily adaptable to different biological contexts and scientific questions.

## SUPPLEMENTARY MATERIALS

**Extended Data Figure 1.**
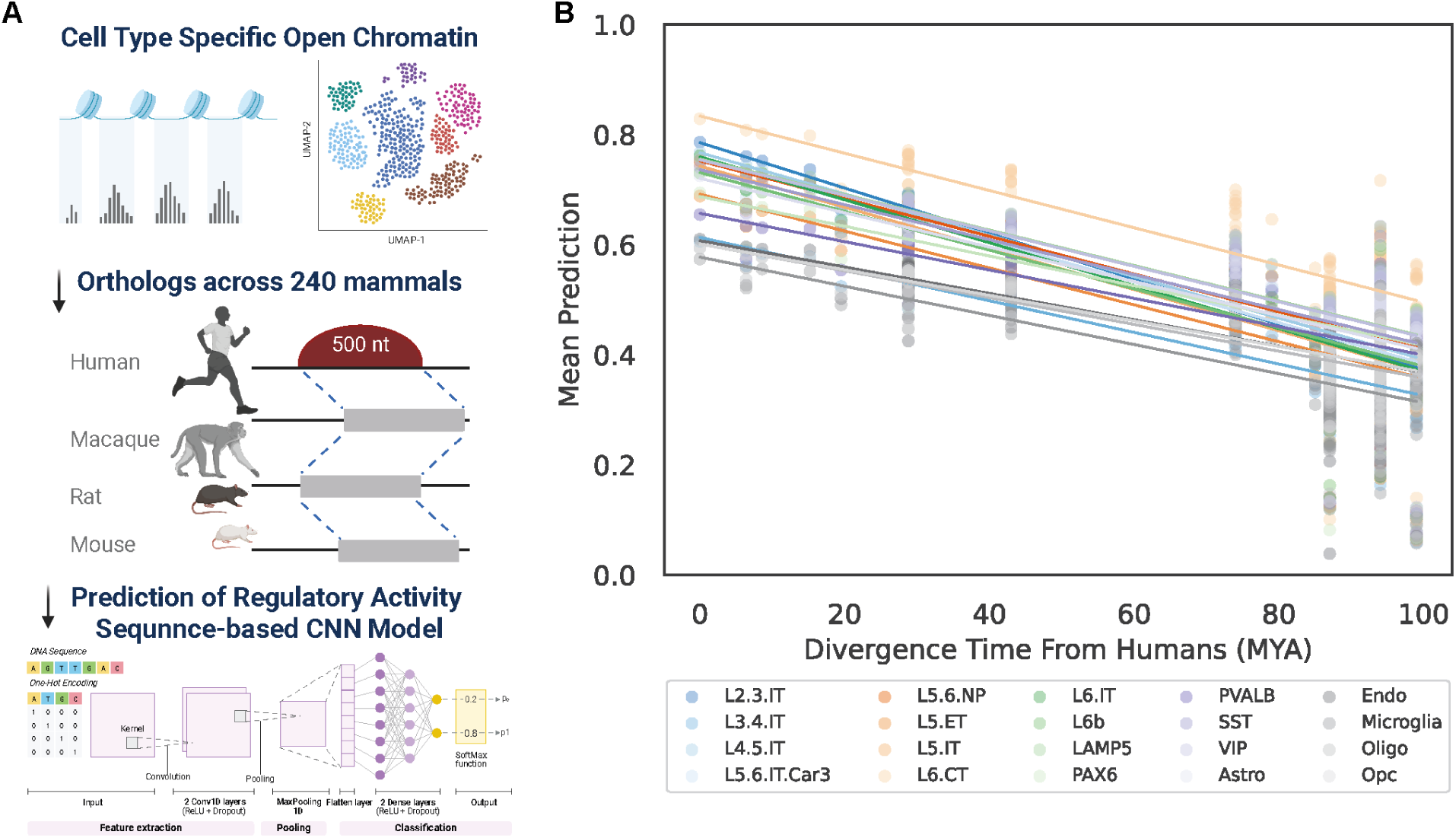
Cell-Type-Aware Conservation Inference Framework. A. Schematic diagram of Sequence-based CNN model workflow. B. Mean predicted enhancer activity across species as a function of divergence time from humans for 20 cortical cell types.

**Extended Data Figure 2.**
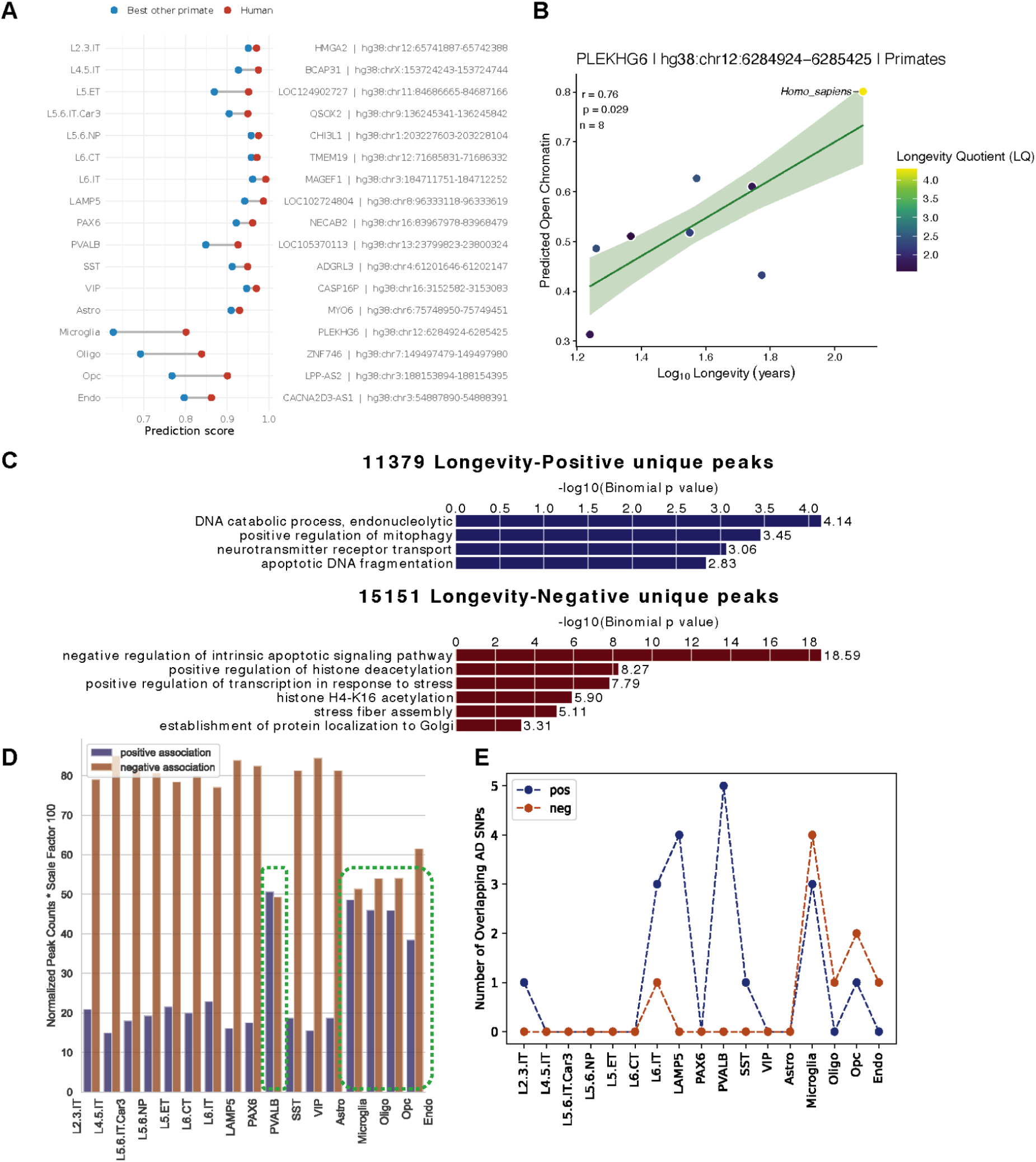
Association of longevity with critical biological processes. A. Most human-specific longevity-associated peak per cell type. For each cell type, top 100 longevity-associated peaks were filtered and ranked by the difference in predicted open chromatin probability in homo sapiens relative to the highest predicted probability in the best other primate in 28 primates. B. Predicted open chromatin regressed on Log Longevity regressed for primates from the Zoonomia^9^ Database. Each point represents one species colored based on the gradient of the longevity quotient. C. Gene ontology enrichment analysis for chromatin regions associated with longevity (positive=top, negative=bottom). Background was set to the whole genome using GREAT. D. Normalized counts of cell-type-specific longevity-associated OCRs (LARs). E. Counts of overlapping AD GWAS genetic variants in these OCRs.

**Extended Data Figure 3.**
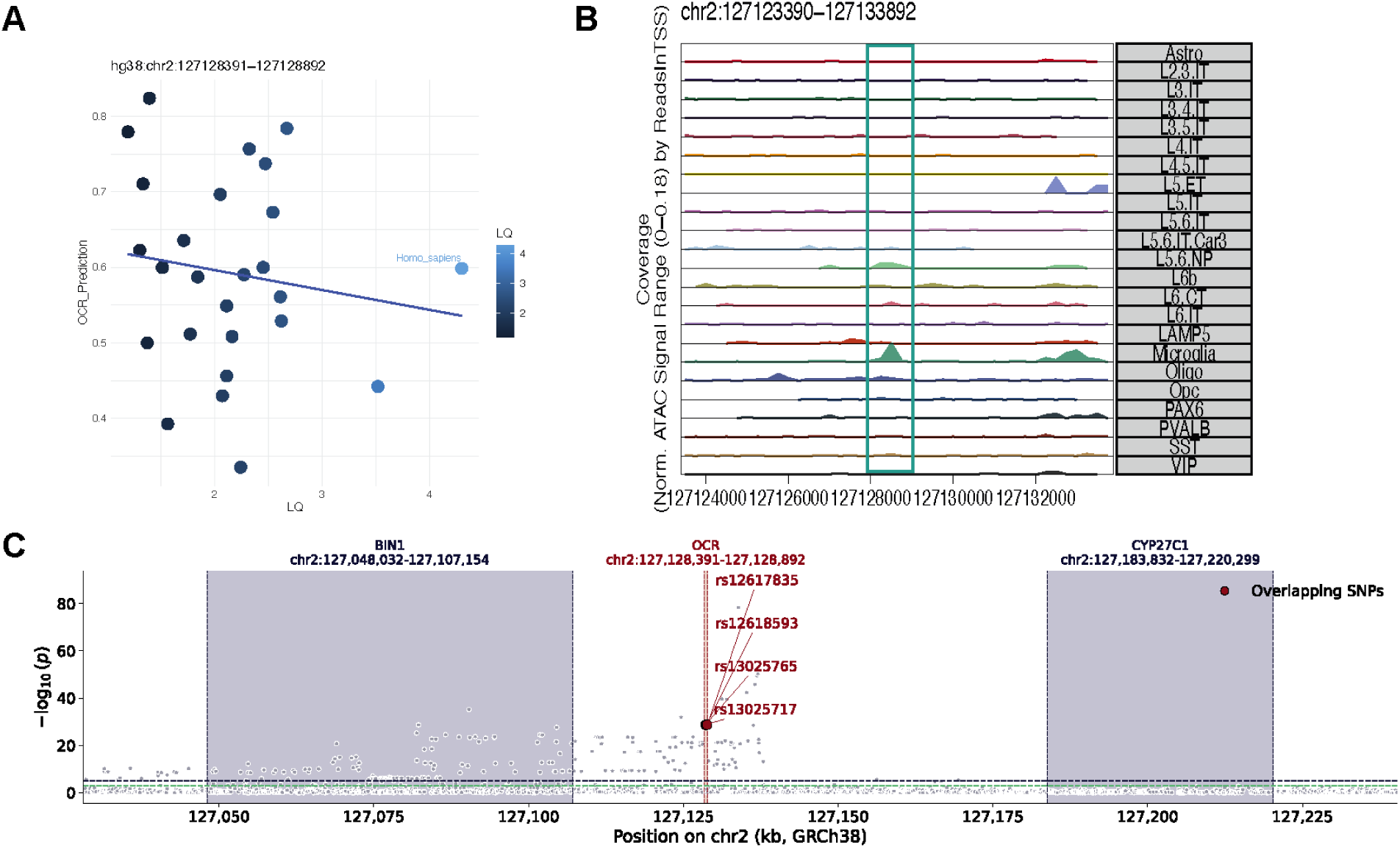
Association of longevity with critical biological processes in Microglia. A. An example of candidate microglia-specific enhancer negatively associated with longevity across primates. B. Browser track plot for microglia-specific enhancer across all cell types. C. Manhattan plot of AD GWAS (−log₁₀ p), enhancer and variants falling in the microglia-specific enhancer are highlighted in red.

**Extended Data Figure 4.**
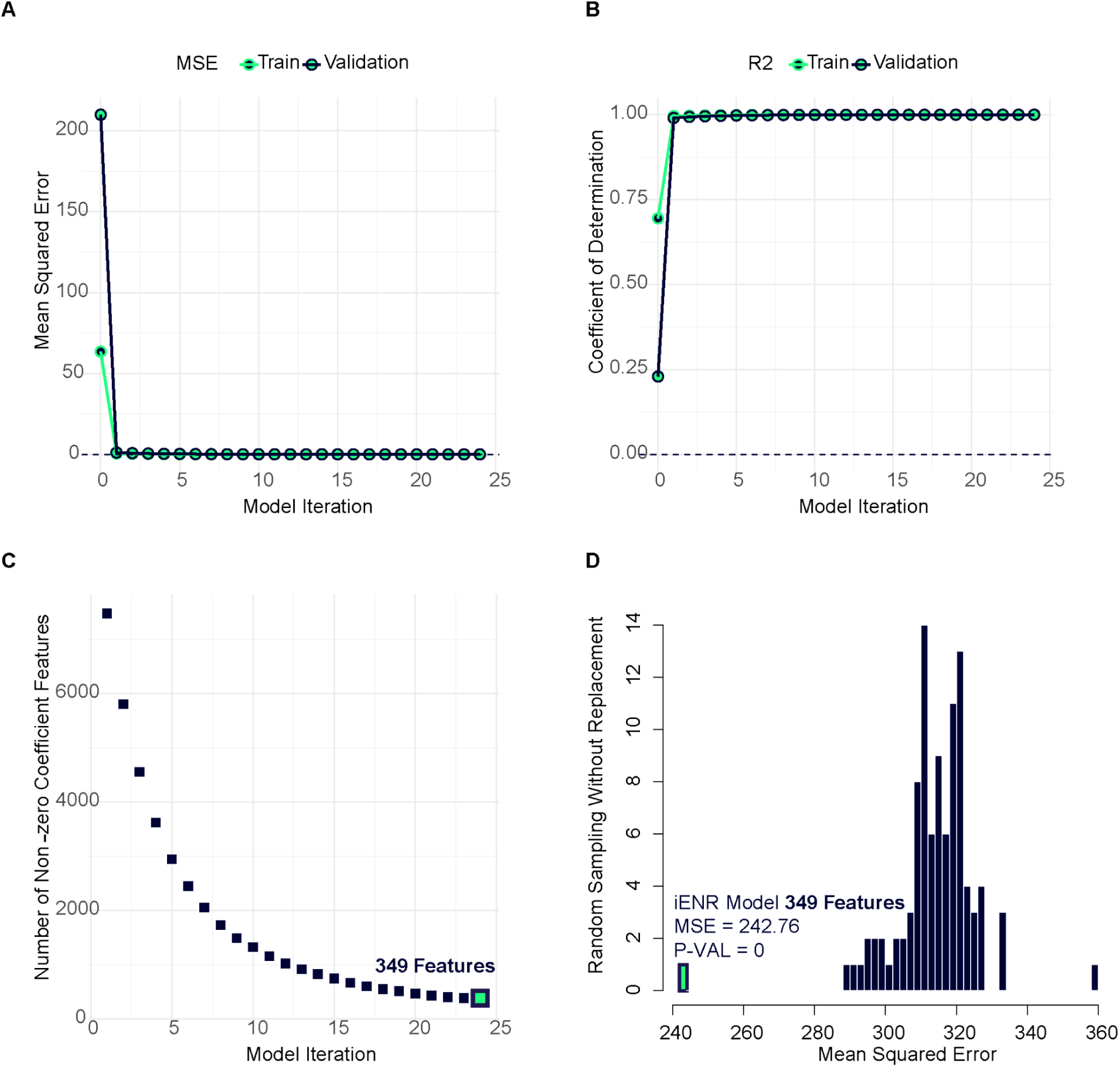
Model performance and feature selection across iterations. A. Mean squared error across 25 iterations for train and validation sets from Ruzicka et al.^20^ dataset comprising healthy controls with age range 24-94 years. B. Coefficient of determination (R^2^) for train and validation sets across 25 iterations. C. Number of fileted critical features across 25 iterations, with the 25th iteration converging on 349 features. D. Mean squared error of predicted cellular age for the validation set using model refined features (349 genes) or an equal set of randomly sampled features without replacement (n=99) demonstrating superior performance of model’s selected features (MSE=242.76, p-value=0).

**Extended Data Figure 5.**
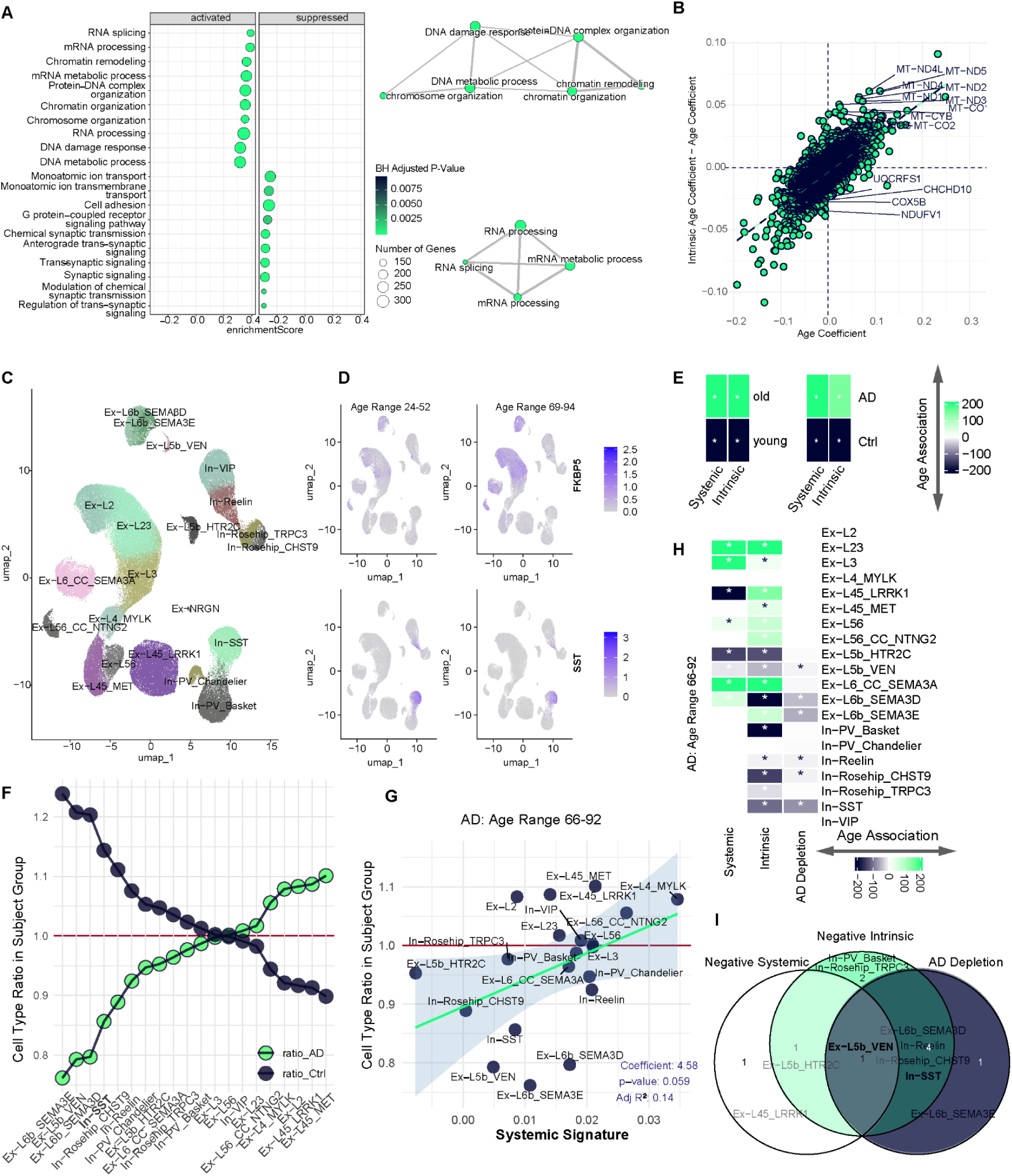
Cell type composition and genetic signatures overrepresentation analysis. A. Gene set enrichment analysis (GSEA) dot plot and enrichment map for genes associated with age as a predictor variable, ranked by the age coefficient. B. Scatter plot of age coefficient (x-axis) and difference of intrinsic age coefficient and age coefficient (y-axis) with 17,658 gene expression features using ordinary least squares linear (OLS) regression model, highlighting mitochondrial- and nuclear-encoded oxidative phosphorylation (OXPHOS) genes. C. Low dimensionality projection of cortical neuronal cell types from Ruzicka et al.^20^ dataset comprising healthy controls with age range 24-94 years. D. Low dimensionality projection feature plot of *FKBP5* (top) and *SST* (bottom) in young subjects 24-52 and older subjects 69-94. Color scale is uniform per feature across conditions, where a feature’s expression is scaled to its maximum expression across the conditions. E. Heatmap for the overrepresentation of cells enriched for aging gene signatures in older vs. younger individuals (left) and AD patients vs. age-matched controls. The color scale represents –log₁₀(*p*) from the hypergeometric test, multiplied by 1 or −1 for gene sets positively or negatively associated with systemic and intrinsic aging. *P*-values were adjusted for multiple comparisons using false discovery rate (FDR) correction applied across all cell types (rows) and features/conditions (columns); adjusted *p*-values are displayed on the plots. F. Ratio of each cell type relative to other cell types in AD patients and aged-matched controls. G. Relation of the mean systemic aging signature activity score per cell type (x-axis) to the ratio of a cell type in AD patients with age range 66-92 years (y-axis). H. Heatmap for the overrepresentation of cells enriched for aging gene signatures per cell type and underrepresentation (depletion) of a cell type in AD subjects. Color scale indicates –log₁₀(*p*) for the hypergeometric test multiplied by +1 for gene sets positively associated with systemic or intrinsic age and –1 for gene sets negatively associated with age. For cell types underrepresented in AD, the –log₁₀(*p*) is multiplied by –1 to reflect depletion. *P*-values were adjusted for multiple comparisons using false discovery rate (FDR) correction applied across all cell types (rows) and features/conditions (columns); adjusted *p*-values are displayed on the plots. I. Venn Diagram for cell types enriched for signatures negatively associated with systemic or intrinsic age and depleted during AD based on hypergeometric tests, highlighting Ex-L5b-VEN to be depleted in AD and enriched for negative systemic and intrinsic signatures.

**Extended Data Figure 6.**
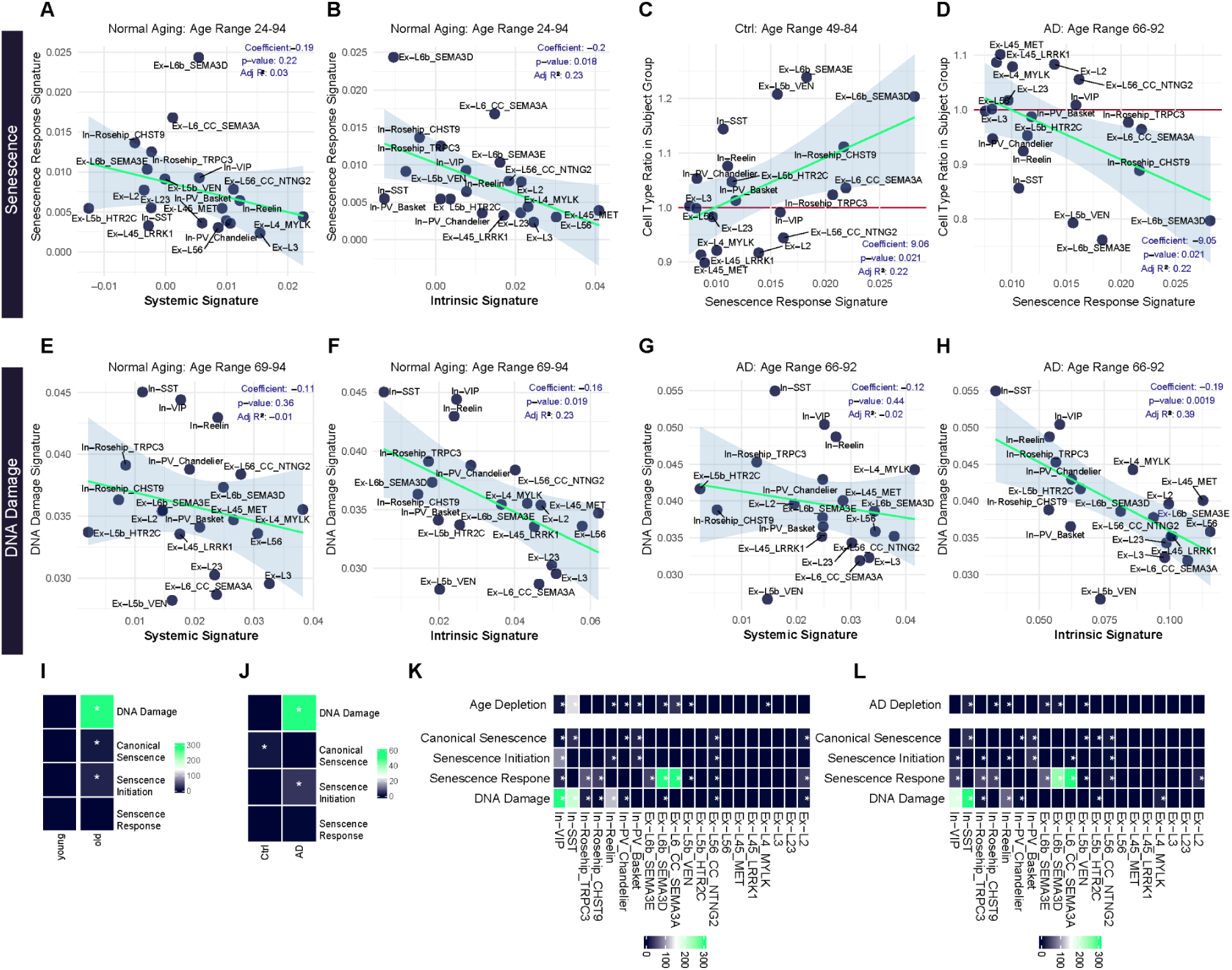
Cell-Type-Specific relationships between systemic and intrinsic aging, senescence programs, and DNA damage in healthy aging and Alzheimer’s disease. A-B. Relation of the mean systemic (A) or intrinsic (B) aging signature activity score (x-axis) and the mean senescence response signature activity score (y-axis) and per cell type in the full cohort of healthy subjects with age range 24-94. In A-H, each point represents a cell type. Linear regression was used to assess associations; regression coefficients, *p*-values, and adjusted R² are displayed on the plots. C-D. Relation of the mean senescence response signature activity score (x-axis) per cell type and ratio of a cell type (y-axis) in healthy aged subjects (C) or in AD subjects (D). E-F. Relation of the mean systemic (E) or intrinsic (F) aging signature activity score (x-axis) and the mean DNA damage signature activity score (y-axis) per cell type in healthy aged subjects. G-H. Relation of the mean systemic (G) or intrinsic (H) aging signature activity score (x-axis) and the mean DNA damage signature activity score (y-axis) per cell type in AD subjects, highlighting the highest DNA damage score for In-SST more than expected based on its cellular age. I-J. Heatmap for the overrepresentation of cells enriched for canonical senescence, senescence initiation, senescence response and DNA damage signatures in older vs. younger individuals (I) or in AD patients vs. age-matched controls (J), highlighting DNA damage to be strongly associated with age and AD. Color scale indicates –log₁₀(*p*) for the hypergeometric test. *P*-values were adjusted for multiple comparisons using false discovery rate (FDR) correction applied across all cell types (rows) and features/conditions (columns); adjusted *p*-values are displayed on the plots. K-L. Heatmap for the overrepresentation of cells enriched for positively or negatively associated aging signatures, canonical senescence, senescence initiation, senescence response and DNA damage signatures per cell type and underrepresentation of a cell type in old subjects (K) or in AD patients (L). Color scale indicates –log₁₀(*p*) for the hypergeometric test. *P*-values were adjusted for multiple comparisons using false discovery rate (FDR) correction applied across all cell types (rows) and features/conditions (columns); adjusted *p*-values are displayed on the plots.

**Extended Data Figure 7.**
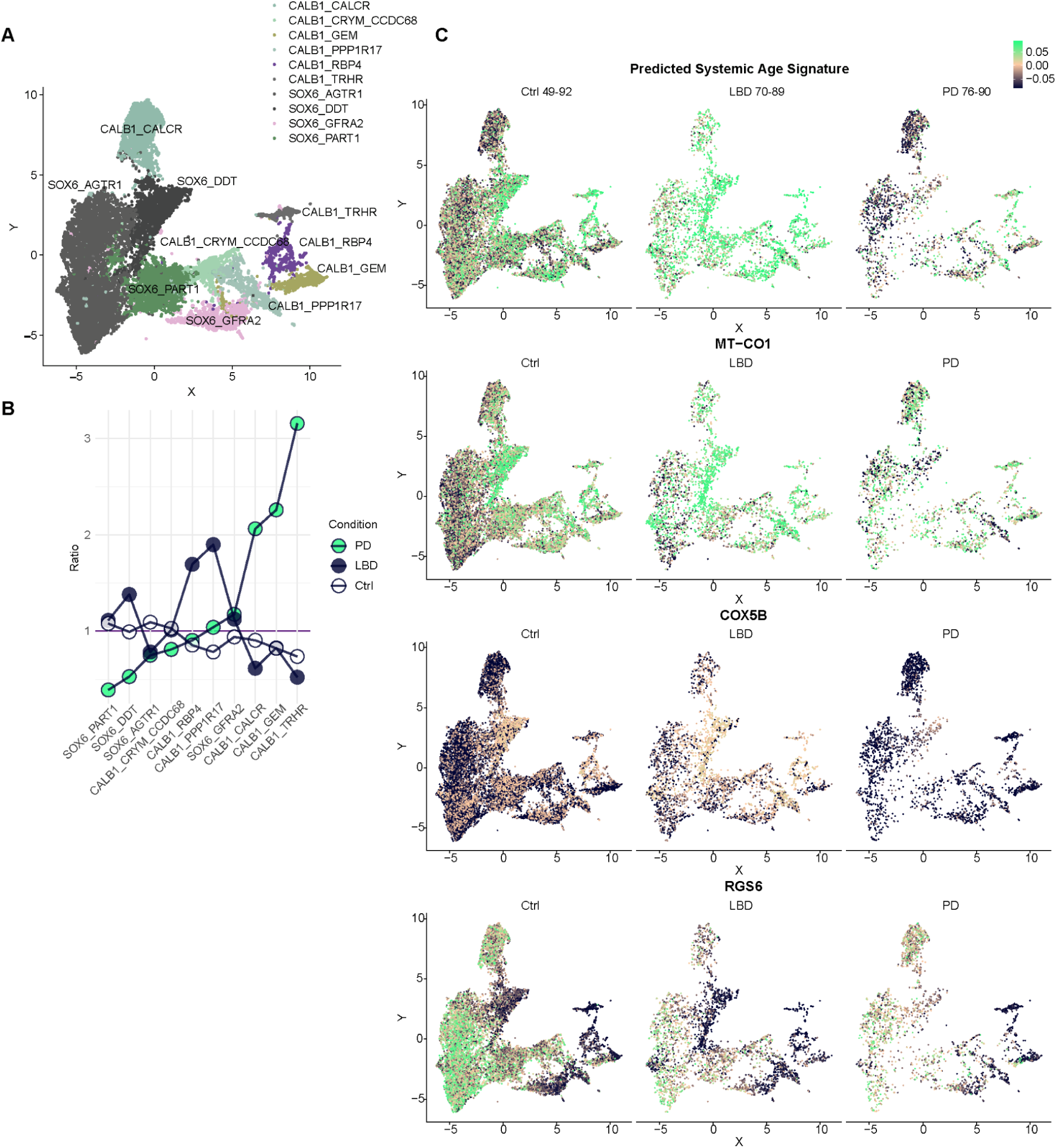
Cell-type composition and aging-associated features in SNpc dopaminergic neurons across Parkinson’s disease and Lewy body dementia. A. Low dimensionality projection of SNpc dopaminergic neurons cell types from Kamath et al.^38^ dataset comprising Parkinson’s disease (PD), Lewy body dementia (LBD) patients and age-matched controls with age range 49-90.^38^ B. Ratio of each cell type relative to all other cell types in PD, LBD patients or age-matched controls. C. Low dimensionality projection feature plot of systemic age (top), MT-CO1 and COX5B (middle), and RGS6 (bottom) in PD, LBD patients and age-matched controls. Color scale is uniform across all plots by scaling plots for all features and conditions to the maximum expression value for the feature with the highest overall expression.

**Extended Data Figure 8.**
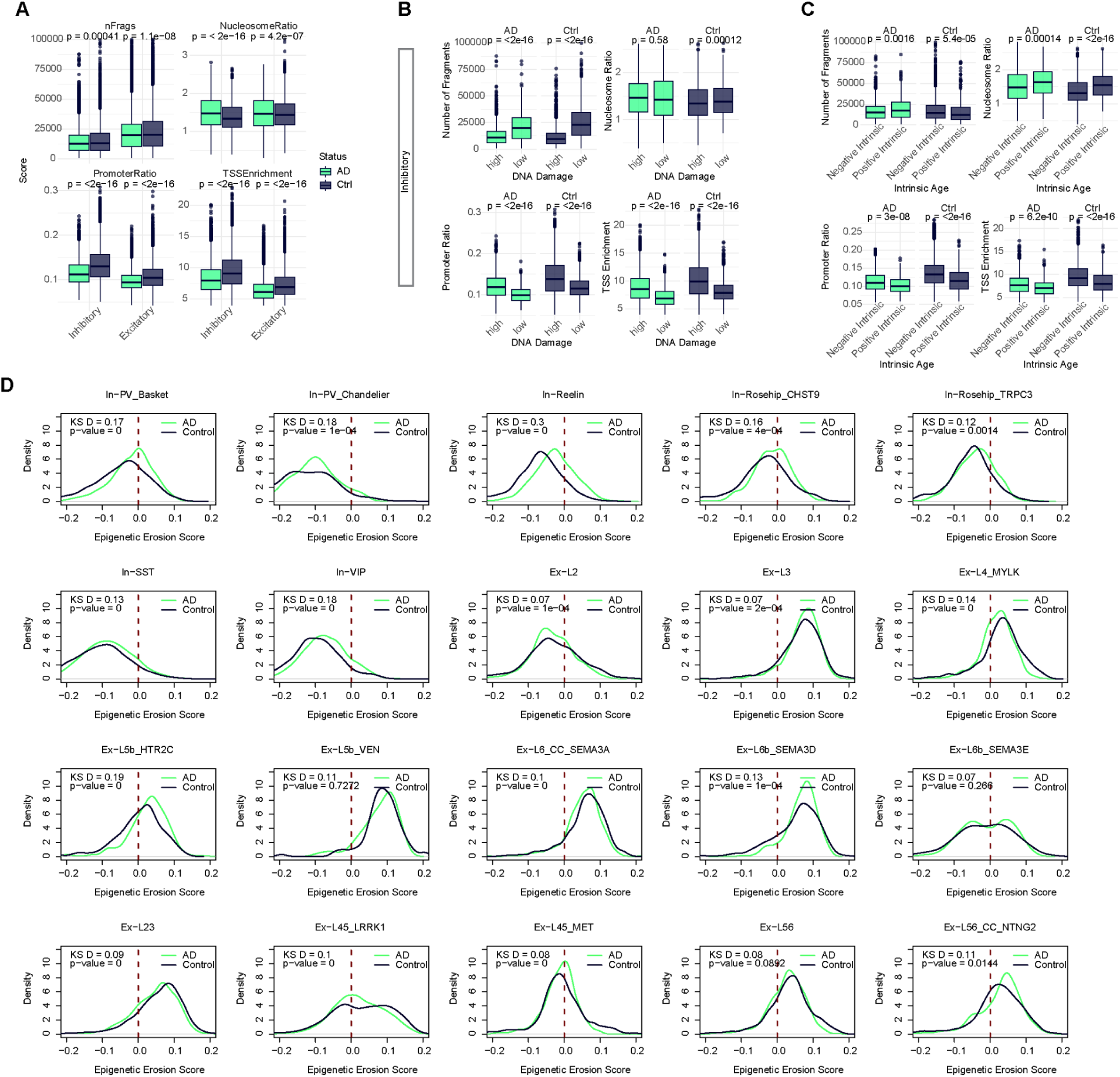
Neuron-type- and signature-specific chromatin accessibility metrics. A. Boxplots for nucleosome ratio, promoter ratio, TSS enrichment and number of fragments split by excitatory and inhibitory neurons in AD patients relative to age-matched controls. Distributions are shown separately for each cell type and subject group. Center lines indicate medians, boxes represent interquartile ranges, and whiskers extend to 1.5×IQR. Statistical comparisons between AD and control groups were performed independently within each cell type using two-sided Wilcoxon rank-sum tests (Mann–Whitney U tests; unpaired, non-parametric). Each observation represents a single cell. Exact raw *p*-values are displayed on the plots. B. Boxplots for nucleosome ratio, promoter ratio, TSS enrichment and number of fragments split by high and low DNA damage signature enriched inhibitory cells in AD patients relative to age-matched controls. C. Boxplots for nucleosome ratio, promoter ratio, TSS enrichment and number of fragments split by intrinsic positive and negative aging signature enriched inhibitory cells in AD patients relative to age-matched controls. D. Density distributions of epigenetic erosion score for neuronal cell subtypes in AD patients relative to age-matched controls. Curves represent kernel density estimates of single-cell epigenetic erosion scores for each group. Distributional differences between AD and control cells were assessed within each cell type using two-sample Kolmogorov–Smirnov tests (two-sided, non-parametric). Each observation represents an individual cell. KS statistics (D) and exact raw p-values are shown on the plots. Red dashed line marks zero.

**Extended Data Figure 9.**
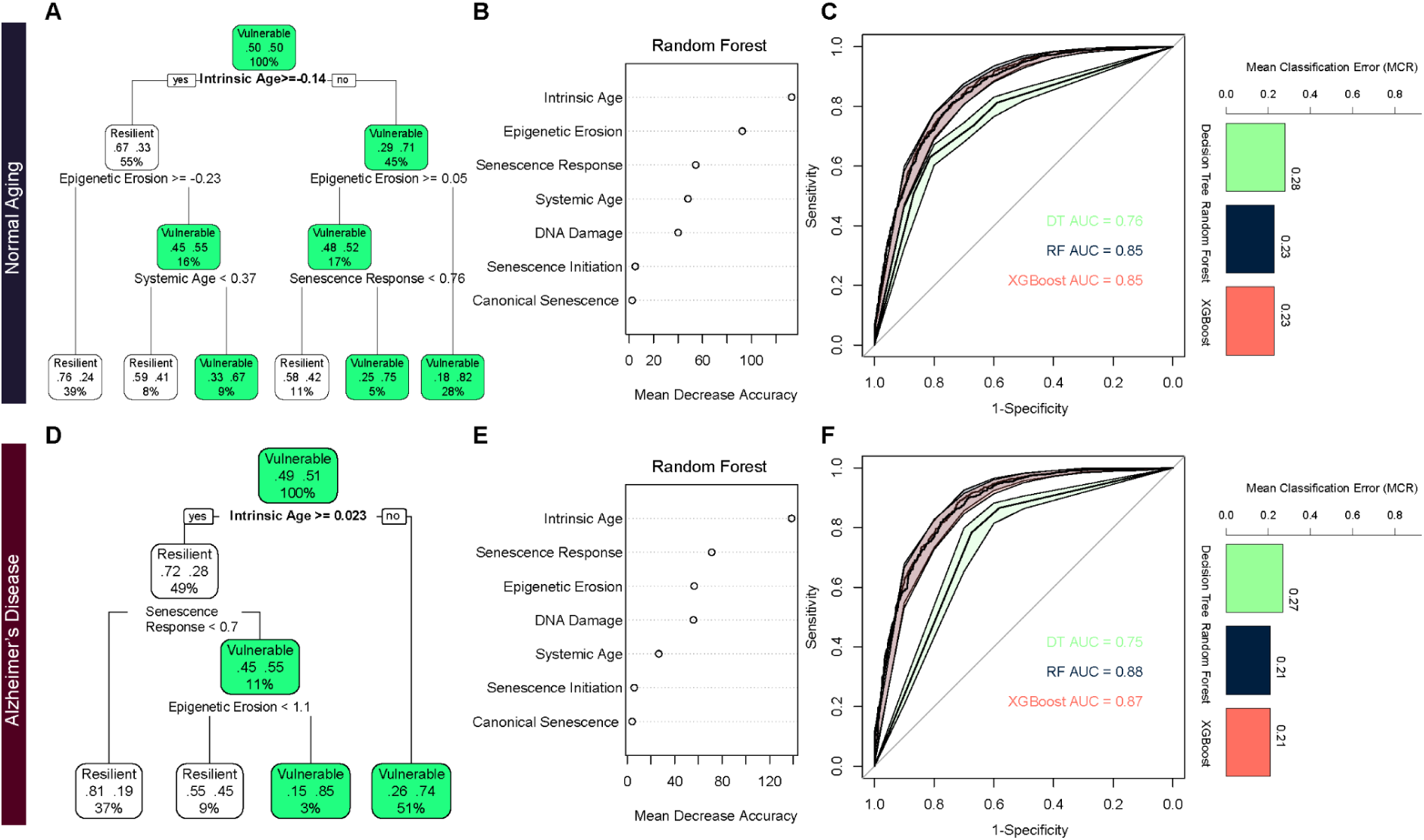
Phenotype-signature-based machine learning classifiers for vulnerability prediction. A,D. Decision Tree classifier was trained to predict vulnerability status (vulnerable vs resilient defined based on cell type composition analysis) in old subjects (A) or in AD patients (D), based on activity scores for signatures of 7 aging phenotypes. Tree complexity was optimized using the 1-SE rule based on cross-validated error, and the final pruned tree is shown. Node colors indicate predicted class, and terminal nodes display class probabilities. B,E. Random Forest classifier was trained using the same feature set. Variable importance scores (mean decrease in accuracy) are shown in old subjects (B) or in AD patients (E), highlighting intrinsic age contributing most strongly to classification performance. C,F. ROC (Receiver Operating Characteristic) curve (1-Specificity vs. Sensitivity) with the area under the curve (AUC) overlaid for direct comparison, with shaded regions indicating bootstrap confidence intervals. Barplots display the misclassification error rate for the three classifiers in old subjects (C) or in AD patients (F). * Models were trained on a designated training set and evaluated on an independent test set; performance metrics were computed on the test set only.

**Extended Data Figure 10.**
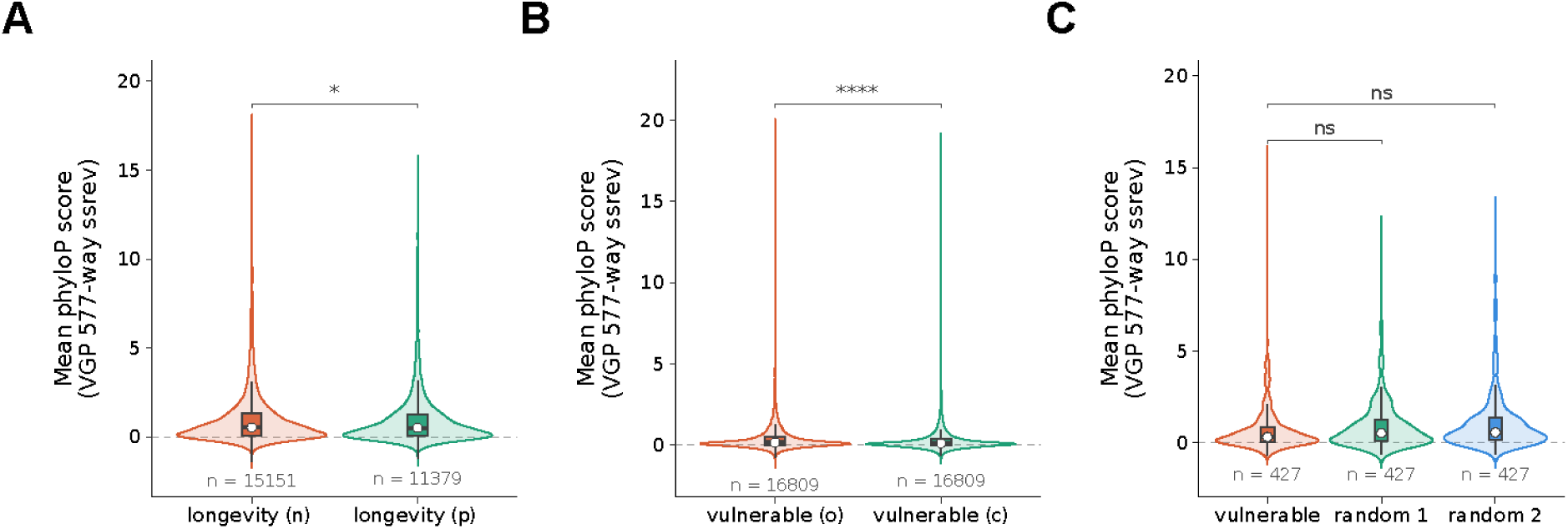
Evolutionary Conservation at the Nucleotide Across Vertebrates. A-C. Distribution of mean phyloP conservation scores (VGP 577-way) for regions associated with vulnerability (o: open, c: closed) (**A**), longevity (n: negative, p: positive) (**B**), VLRs relative to two randomly sampled sets (**C**). Violins show the full distribution with embedded boxplots showing the interquartile range with the median marked as a white point. The dashed line marks phyloP = 0. Sample sizes (*n*) are indicated below each group. Significance was assessed by a one-sided Mann-Whitney *U* test.

**Supplementary Tables 1-6**

- **Supplementary Table 1.** Open chromatin regions positively-associated with longevity (11,379 OCRs, BH-corrected p-value < 0.05).
- **Supplementary Table 2.** Open chromatin regions negatively-associated with longevity (5,151 OCRs, BH-corrected p-value < 0.05).
- **Supplementary Table 3.** Systemic age signature from ENR model with each gene’s coefficient.
- **Supplementary Table 4.** Intrinsic age signature from iENR model with each gene’s coefficient.
- **Supplementary Table 5.** Differentially accessible regions in vulnerable neurons (inhibitory neurons enriched in negative intrinsic age signature) in Alzheimer’s disease (3,362 vulnerability-associated regions (VARs)).
- **Supplementary Table 6.** Shared regions between mammalian longevity and human age vulnerability (427 vulnerability and longevity regions (VLRs)).

**Supplementary Information**

## METHODS

### Data Processing for Human Prefrontal Cortex snRNA-Seq

Single-nucleus RNA-Seq (snRNA-Seq) data from the human prefrontal cortex neurons (PFC), were obtained from were obtained from Ruzicka et al. (2024, Science), comprising 69 controls with age range 24 to 94 years old, 189,423 nuclei and 17,658 genes across 14 excitatory cell types and 7 inhibitory annotated cell types ^20^.

The original study employed a sample multiplexing strategy, pooling cases and controls within each sequencing library to limit batch-driven confounding during library preparation. The 58 control individuals used here were distributed across 24 batches (2-4 individuals per batch).

To conserve biological variation relevant to cellular age prediction — as aggressive batch correction has been shown to attenuate genuine transcriptomic signals ^66^ — we proceeded with standard normalization and clustering without additional batch integration, in line with the processing approach of the original study. Data was processed using Seurat NormalizeData() and ScaleData() functions in Seurat v5.

For model training, differences in sequencing depth between individuals were addressed by including the per-individual mean UMI count (nCount_RNA_ID) as a covariate during model training. Post-hoc donor-level analysis confirmed that PMI showed no association with residuals from a linear model of predicted intrinsic age on chronological age (Pearson’s r = 0.001, p = 0.992); sequencing batch showed a non-significant association with the same residuals (one-way ANOVA, p = 0.06).

Data were partitioned at the donor level, stratified by age group and sex, into training (n = 49, 133,323 nuclei), validation (n = 9, 29,342 nuclei), and test (n = 11, 26,758 nuclei) sets. The validation set was used for model selection during hyperparameter optimization; the test set was held out until final evaluation. Once the optimal model architecture was finalized, the test dataset was incorporated for downstream analyses. To avoid information leakage across cells from the same individual, cells were restricted to individuals assigned to the training and validation sets. Gene expression values were log-normalized using Seurat package in R. Features and covariates were subsequently standardized using a StandardScaler fit exclusively on the training data and applied to the validation and test sets using scikit-learn in Python.

For external validation and AD-context evaluation, we used human dorsolateral prefrontal cortex (DLPFC) single-nucleus multiomics (snRNA-Seq and snATAC-Seq) dataset comprising 105,332 nuclei isolated from 7 AD and 8 unaffected donors with age range 49 to 92 ^38^. Pre-processing included removal of ambient RNA by SoupX ^67^ and doublet detection using DoubletFinder ^68^. Data was processed using Seurat NormalizeData() and ScaleData() functions in Seurat v5 and cell type labels were transferred from the training dataset ^20^. For PD/LBD-context evaluation, we utilized human substantia nigra pars compacta (SNpc) snRNA-Seq dataset comprising 22,048 dopaminergic neurons across six PD (age range 76-90), four LBD donors (age range 70-89) and eight neurotypical donors (age range 49-92) ^40^.

### Data Processing for Human Prefrontal Cortex snATAC-Seq

Using the DLPFC dataset^38^, we adapted the implementation for the ArchR package ^69^. The pipeline involved creating ArrowFiles, inferring and filtering doublets and creating an ArchR project for downstream analyses. With peak calling with Macs2, reproducible peaks are identified across pseudobulk replicates for each group in the variable of interest. We identified chromatin accessibility profiles for major cell classes and neuronal subtypes separately. This process involved: (1) generating of cell type x sample pseudo-bulk replicates using addGroupCoverages function (minCells = 40, maxCells = 2000, minReplicates = 2, maxReplicates = 12), and (2) calling peaks with MAC2 ^70^ and constructing a union reproducible peak set using addReproduciblePeakSet function (reproducibility = "(n+1)/2", genomeSize = 2.7e9, peaksPerCell = 500, maxPeaks = 300000, minCells = 25, cutOff = 0.1, promoterRegion = c(5000, 100)). This yielded a cell x peak matrix through ArchR.

### Peak Calling and Mapping of Orthologous Regions Across Species

We identified open chromatin regions from cortical snATAC-Seq data in four species: humans^22,23^, rhesus macaques^24^, rats^25,26^ and mice^27–29^. To map orthologs from 4 source species to 240 mammalian genomes from the Zoonomia database^9^, we leveraged CACTUS whole-genome multiple sequence alignments^71^ with haLiftover^72^ to identify putative orthologs of genomic regions from query species in a target species. For postprocessing, we used halLiftover Post-processing for the Evolution of Regulatory Elements (HALPER)^73^, for generating contiguous putative region orthologs from the outputs of halLiftover.

### Cell-TACIT Model

We trained Cell Type-Aware Conservation Inference Toolkit (Cell-TACIT) convolutional neural network models (CNN) in the cortex using cell type OCRs derived from our scATAC-seq experiments. Our inputs were one-hot encoded, 500 base pair DNA sequences, and the outputs varied based on whether we trained a classification or regression model: classification model outputs were a probability within [0,1] that the input OCR was accessible, and regression model outputs were a continuous-valued number greater than or equal to zero, representing the predicted strength of the ATAC-seq peak signal. All models used positive sets from the reproducible OCR set detected in their cell types across species, where we removed OCRs overlapping exons and within 20,000bp from a TSS to thoroughly exclude promoter elements, because promoters and enhancers within the same cell type have been shown to be bound by only partially overlapping groups of transcription factors.

We trained 5-fold cross-validated models using the following chromosome hold out scheme: OCRs on the test set (chr1 and chr2) are never seen by any models or evaluations for hyper-parameter selection, and models for each fold do not see OCRs on its own validation chromosome hold-out sets: fold1: chr[6, 13, 21], fold2: chr[7, 14, 18], fold3: chr[11, 17, X], fold4: chr[9, 12], and fold5: chr[8, 10]. In the cross-species setting, the peak assignment to test, validation, and training sets are determined by the hg38 chromosome. Mappable peaks were assigned by their own hg38 chromosome, unmappable peaks were assigned by the hg38 chromosome of the mapped peaks in synteny with them. To identify peaks in synteny with an unmappable peak, we sorted the peaks by the source (mouse or macaque) chromosome and coordinates and converted the results to a data table. We then filled in the missing hg38 chromosome –corresponding to a peak not mappable to hg38– by the previous or next non-missing hg38 entry by running dplyr::fill(direction = “updown”).

### Classification Cell-TACIT Models

We trained classification models using the following negative sets to identify the negative set with the best balance in overall performance and specific performance at detecting cell type OCR specificity and changes in OCR conservation.

1. Non-Enhancer Orthologs Negatives: We obtained orthologs of the positive set OCRs mapped to other species that did not overlap any reproducible or non-reproducible OCR of the cell type used in model training. This negative set aims to push models to learn cases when cell type OCR activity is lost although genome sequences can still be aligned.
2. GC-Matched negatives: We obtained the large set of GC-matched negatives to the positive set within each species using the BiasAway tool^74^ with the following parameters: biasaway c --nfold 10 --deviation 2.6 --step 50 --seed 1 --winlen 100. While 10 times GC-matched negatives are generated, we further filter them to exclude exonic and promoter regions and regions overlapping any reproducible or non-reproducible OCR of the target cell type, resulting in 4-5 times as many GC-matched negatives as positives. This negative set aims to push models to learn to identify random non-coding regions in the genome that are inactive, but have a similar GC-content to our training set.
3. Differential Cell Type Negatives: We identified differential OCRs by intersecting the orthologs of the positive set OCRs with OCRs from a second cell type in the same species that were not active in the training cell type. This was done using ArchR, where we first created pseudo-bulk replicates per cell type using addGroupCoverages() and identified peaks using addReproduciblePeakSet(). We then quantified peak accessibility across cell types using getPeakMatrix() followed by getMarkerFeatures() with bias = c("TSSEnrichment", "log10(nFrags)") to control for technical covariates. We defined a peak as differential if it had a log2 fold change ≥ 1 and FDR ≤ 0.01 in the second cell type compared to the training cell type. From these, we extracted peaks that were significantly more accessible in the second cell type but not present in the training cell type’s reproducible or non-reproducible peak set, using getPeakSet() and custom filtering with GenomicRanges::findOverlaps() and subsetByOverlaps(). We then intersected these cell-type-specific peaks with the orthologs of the positive OCRs and further filtered to exclude any that overlapped OCRs in the training cell type. This negative set aims to push models to learn to distinguish regions that are active in other cellular contexts but inactive in the target cell type, encouraging learning of cell-type-specific chromatin accessibility patterns despite sequence conservation. While the original TACIT method uses OCRs that are open in all other cell types other than the foreground, we chose to use our differentially expressed OCRs to allow the models to learn a more nuanced pattern of accessibility across different groups of cell type backgrounds.

To develop classification models of chromatin accessibility, we use a CNN architecture with cyclic learning rate (LR) and momentum to accelerate training time. For hyperparameter optimization, we swept over the initial and max cyclic LR, width/number of the convolutional layers, the number of convolutional filters, and the convolutional layer dropout rate. We swept over all of the cell type models separately, and found that there was virtually no difference in performance comparing cell type-specific hyperparameters vs using a uniform set. As such, we set an initial/max LR of 0.002-0.49, 0.87-0.99 for the min/max momentum, 6 convolutional layers 13 units wide, 500 convolutional filters, and a dropout rate of 0.3 for all of our models. Following the initial training for 24 epochs, these models were then fine-tuned for a further 10 epochs, exponentially decaying our learning rate at a rate of 0.95. All models were trained using a proportional loss function and stochastic gradient descent, with a batch size of 100.

### Cell-TACIT Model Performance Evaluation

We evaluated Cell-TACIT models across folds for overall performance on the validation set using the area under the receiver-operator curve (AUROC) and area under the Precision-Recall curve (AUPRC). To assess our Cell-TACIT models’ ability to detect cell-type specific differences in open chromatin, we constructed evaluation sets consisting of OCRs accessible in only the target cell type and OCRs only accessible in all other cell types. To assess the ability of our Cell-TACIT models to detect species-specific open chromatin changes, we predict the activity state in cell type OCRs where the ortholog in another species is accessible or where the ortholog is inaccessible. Lastly, we accessed the average Cell-TACIT predictions of cell type OCRs when mapped to increasing distant species. This was done to assess the phylogeny-matched correlations by determining the relationship between the model predictions and the years diverged from source OCRs.

When evaluating the models on the general validation set, we observed that the models’ AUROC ranged from 0.81-0.87 over the glial cell type models, and 0.79-0.9 for the neuronal cell type models. The validation AUPRC ranged from 0.5-0.71 over the glial cell type models, and 0.56-0.82 for the neuronal cell type models. Given this range of performance, we also assessed the correlation between the validation performance and the amount of training data available. We observed a Pearson correlation coefficient of 0.44 for the AUROC metric/number of training peaks, and a coefficient of 0.52 for the AUPRC metric/number of training peaks, indicating a degree of correlation between our model performance and the amount of training data available for a given cell type. We also posit that certain cell types may be less biologically distinct, and their comparatively murkier definition may result in the lower performance as compared to more distinct subtypes.

When evaluating on the cell-type specific validation set, the models’ AUROC ranged from 0.58-0.86. When evaluating on the species-specific validation set, the models’ AUROC ranged from 0.63-0.78. This indicates that the models are able to learn both cell type and species-specific patterns of chromatin accessibility when evaluated on our validation set.

We do not expect all species to use the same set of OCRs across species. Furthermore, we expect the OCRs measured in one species will be less active on a genome-wide level when mapped across species in more distant species due to evolutionary drift and usage of cell type-specific gene regulation. We mapped human OCRs across species and observed the expected decline in the number of OCRs that can be mapped in increasingly distant species. This mappability may also be lower in certain species due to lower quality genomes or representation within Zoonomia. From these orthologous DNA regions, we found a negative correlation with the average validation predictions of each species and the number of years diverged from humans. This phylogeny-matched correlation analysis suggests our models were able to predict the regulatory decay of cell type OCRs across species.

Furthermore, down-sampling analysis was performed to determine the least number of peaks that can be used to train an accurate CNN model and achieve high performance (AUROC > 0.8 and AUPRC > 0.65). The minimum number of peaks was ≥ 8000 peaks per cell type. We do not expect difference across species or gender to affect the predictions derived from the model. As the influence of cell type-specificity on the variation of open chromatin exceeds the influences of other inter- and intra-species variability, such stress, age, and sex. Additionally, similar proportional representation of both sexes for each species have been included.

### Cell-TACIT Predictions and Construction of Mammalian Cell-Type-Specific Chromatin Accessibility Atlas

We predicted open chromatin at orthologs of human cortical cell type OCRs that were mappable to each Zoonomia genome. This creates a peak-by-species matrix for each cell type with Cell-TACIT predictions, where there is a row for each human OCR for a cell type and a column for each Zoonomia genome. This allowed us to construct an atlas of experimentally measured and predicted accessible chromatin across 18 cell types and 240 mammalian genomes. This atlas comprised **3,180,197** peaks measured in four reference species. The experimentally measured peaks were then mapped to orthologous regions across 240 mammalian species, yielding a **3,180,197 × 240 matrix** and a total of **763,247,280** measured and predicted accessible chromatin regions. This comprehensive atlas provides a resource for exploring cell-type-specific chromatin accessibility across the mammalian lineage.

Breakdown of the number of OCRs by cell-type:

- **Excitatory neurons:** L2.3.IT (322,945), L4.5.IT (285,812), L5.ET (142,084), L5.6.IT.Car3 (192,027), L5.6.NP (132,211), L6.CT (125,500), L6.IT (292,635) — total 1,493,214 peaks.
- **Inhibitory neurons:** LAMP5 (136,731), PAX6 (140,696), PVALB (219,528), SST (228,304), VIP (155,672) — total 880,931 peaks.
- Glia: Astrocytes (205,151), Microglia (117,914), Oligodendrocyte precursor cells/OPCs (183,245), Oligodendrocytes (160,452) — total 666,762 peaks. Vascular cells: Endothelia (42,810), Vascular leptomeningeal cells/VLMC (96,480) — total 139,290 peaks.

To retrieve divergence time of the mammals from homo sapiens, we used TimeTree using PAReTT ^75^. To evaluate predictions of a species-specific model, we regressed model predictions across 240 mammals for 17 cortical cell types on the divergence time. This reflected an inverse relationship between human-specific model predictions and divergence time from humans.

As introduced in Srinivasan, Phan *et al*, we calibrated the raw Cell-TACIT model score by the percentage of the validation set that had that score or lower, in order to scale the strength of CNN nonlinear sigmoid activation across model folds and cell types. This was accomplished by taking the Cell-TACIT score of the validation positive set to construct an empirical cumulative distribution (ecdf R function), and then scaled all new predictions by the fraction of validation positives with a lower score. This produced calibrated scores near 0 for the validation negative set and expanded the dynamic range for prediction values in the positive set. We applied this calibration to the score for the average across the species groupings for Cell-TACIT predictions and individual species prediction. For each cell type, we classified whether each human OCR is predicted to be accessible in each of the 20 groupings by defining predicted accessible as having a calibrated prediction of at greater than 0.5, yielding binary annotations of accessibility.

### Application of Tissue-Aware Conservation Inference Toolkit (TACIT)

Using the constructed atlas, we applied Tissue-Aware Conservation Inference Toolkit (TACIT) ^10^ which combines phylogenetic linear regression (PhyloLM) and phylogenetic permutations (permulations) to identify OCRs associated with longevity quotient (LQ), controlling for the relationship between lifespan and body size (Figure 2A). Cell type-specific LQ-associated OCRs were derived based on PhyloLM without permutations and were evaluated for their implication in neurodegenerative disorders based on their intersections with SNPs from AD GWAS. Additionally, GREAT was used for ontology enrichment analysis of longevity associated regions, enrichment results were visualized with the assistance of a large language model (Claude Opus 4.8), and all values were verified against the GREAT output (Figure 2B).

### Baseline Systemic Aging Model

We employed an elastic net regression model, which is a form of regularized linear regression that penalizes a model for large feature coefficients. We first trained a standard ElasticNet regression model (iteration 0) to predict chronological age from single-cell gene expression profiles. This model serves as a baseline and captures dominant, tissue-wide (systemic) aging signals shared across cells. The model was implemented using ElasticNetCV (scikit-learn) with 5-fold cross-validation to select optimal regularization parameters:

- Mixing parameter (L1 ratio / λ): controls the relative contribution of L1 (lasso) and L2 (ridge) penalties.
- Regularization strength (α): controls the weighting of the sum of both penalties to the loss function.

The systemic model was fit using gene expression features as the predictor variables and chronological age as the target variable. Cells derived from the same individual are initially labeled with the same starting chronological age. To account for technical variation, we included sequencing depth as a covariate:

- Per cell: reciprocal of total transcript counts per cell (1/nCount_RNA_Cell) as a sample weight.
- Per individual: mean sequencing depth per donor (nCount_RNA_ID) as donor-level sequencing depth covariate (nCount_RNA_ID).

*Age/cell ∼ Features + nCount_RNA_ID*

*Mathematical Equation:* **ŷ^(0)^ = ƒ_EN_(X, C), y^(0)^ = y**

*y: Chronological age

*X: Full set of features

*C: Covariates

Model performance was evaluated on both training and validation sets using: mean squared error (MSE), mean absolute error (MAE), coefficient of determination (R²).

### Iterative Intrinsic Aging Model

To capture cell-specific autonomous aging from the shared aging signal, we extended the baseline model using an iterative refinement framework over 25 iterations following the same architecture of the baseline model. Predicted age from the baseline model (iteration 0) was retained as input for the first iteration of the intrinsic model (iteration 1), where predicted age substitute age as the new target variable and non-zero coefficient features are used as the new filtered set of predictors with chronological age and mean sequencing depth per donor as covariates. The next iteration is initiated, with the latter’s output being the input for the new model for a set number of iterations (i=25).

*Predicted age/cell ∼ Non-zero coefficient features + nCount_RNA/ID + Donor Age*

*Mathematical Equation:* **ŷ^(t)^ = ƒ_EN_(X_S(t-1)_, C), y^(t)^ = ŷ^(t-1)^, for t≥1**

*ŷ^(t-1)^: Predicted cellular age from previous iteration (t-1)

*X_S(t-1)_: Subset of features from previous iteration (t-1)

*C: covariates

Over each iteration, the model learns to identify genes that are related to whether a cell displays accelerated aging relative to other cells within a given subject, rather than genes that are broadly associated with aging. All scaling steps for features and targets were performed using parameters fit on the training data at each iteration and applied to the validation data.

#### Iterative procedure

At each iteration t≥1:

1. **Feature selection** Features with non-zero coefficients from the previous iteration were retained. This progressively refines the feature space toward genes most informative for modeling cell-level variation.
2. **Target update** The predicted age from the previous iteration (ŷ^(t−1)^) was used as the target variable for the current model.
3. **Model fitting** The model was fit using the same cross-validation and weighting scheme as in the baseline model. A new ElasticNet regression model was trained using:

- the subset of selected genes
- sequencing depth covariate (nCount_RNA_ID)
- donor chronological age covariate (included as an additional covariate in later iterations)
4. **Prediction and propagation** Predicted values from the current iteration (ŷ^(t)^) were computed for both training and validation sets and propagated forward as input to the next iteration.

#### Tracking Table for Model Training

| Model | Description | Target | Predictor | Co-variables | Weight |
| --- | --- | --- | --- | --- | --- |
| Iteration 0 | Standard ElasticNet | Donor Age | Full Gene Set | nCount_RNA_ID | 1/nCount_RNA_Cell |
| Iteration 1 | Iterative ElasticNet | Predicted Age $\hat{y}^{(t-1)}$ | Refined Gene Set from t-1 | nCount_RNA_ID + Donor Age | 1/nCount_RNA_Cell |
| Iteration 2 | Iterative ElasticNet | Predicted Age $\hat{y}^{(t-1)}$ | Refined Gene Set from t-1 | nCount_RNA_ID + Donor Age | 1/nCount_RNA_Cell |
| Iteration N | Iterative ElasticNet | Predicted Age $\hat{y}^{(t-1)}$ | Refined Gene Set from t-1 | nCount_RNA_ID + Donor Age | 1/nCount_RNA_Cell |

### Aging Model Performance Evaluation

The baseline (iteration 0) model was evaluated against chronological age, providing a direct benchmark comparable to existing transcriptomic aging clocks. Evaluation metrics included Pearson correlation, MAE, and MSE.

For iterative models (iteration > 0), performance was evaluated against both:

- chronological age to assess biological relevance and comparability to standard aging models
- iteratively refined predicted age (ŷ^(t−1)^) to assess internal consistency and stability

### Gene Selection and Signature Definition

By developing an iterative learning approach, information learned from a model feeds into a new model until reaching a model that converges on the most informative features associated with differences in intrinsic aging. This allowed for identifying aging associated signatures that are independent of tissue systemic effects highlighting the disparity in the implication of certain genes to the aging process at different scales. The final intrinsic aging signature was defined as the set of genes with non-zero coefficients in the final iteration. This iterative feature selection process progressively reduced the feature space from the full transcriptome (17,658 genes) to a sparse subset (349 genes), representing genes most strongly associated with cell-specific aging variation.

### Aging Model Interpretability and Signature Evaluation

Comparing elastic net performance on the defined feature set versus equal-sized random sets, the predicted-age signature model outperformed 99 random models, achieving a mean squared error (MSE) of 242.8 versus 288.1 for the best random model. The final model retained 349 unique features, a more stringent set than the first iteration. Applied to the AD dataset, the intrinsic signature was more heterogeneous across cell types than the systemic signature, demonstrating that our approach effectively disentangled cell type-specific from systemic effects.

### Quantification of Gene Signature Activity Scores

The model parsed out systemic and intrinsic aging genetic signatures. This allowed for identifying cells actively expressing these signatures by assessing gene set activity in each cell using AUCell^50^ on snRNA-Seq data from the human prefrontal cortex of healthy individuals ^20^ and AD patients ^38^, as well as, SNpc of Parkinson’s disease (PD) and Lewy body dementia (LBD) patients ^40^. This resolved the differential enrichment of the identified signatures in cortical and midbrain cell subtypes. For AUCell, we adapted the implementation for the R/bioconductor package available at http://scenic.aertslab.org ^76^. Our pipeline involved filtering non-zero coefficient features and binning them based on the direction of the coefficient into positive and negative age associated signatures. This was followed by building gene-expression rankings for each cell and calculating enrichment for the gene signatures (AUC) by setting a threshold to the top 5% genes in the cell ranking and then calculate the area under the curve (AUC) from the recovery curve which reflects the proportion of gene signature in the top 5% genes in a cell. Activity scores were computed for each signature per cell. The assignment of enriched cells was based on the Global k1 threshold, which denotes the global distribution (mean and standard deviations of all cells). The threshold is set to the right of the distribution (prob: 1-(thrP/nCells)), where thrP is the probability to determine outliers (if the AUC is normally distributed) and by default it is set to 1%.

### Gene Set Enrichment Analysis (GSEA)

We regressed the full set of 17,658 features on chronological age using ordinary least squares linear regression model (OLS), while accounting for the mean number of counts per subject. Then, we employed the Gene Set Enrichment Analysis (GSEA) method to the set of features significantly associated with age, either positively or negatively, sorted by the age coefficient. To capture the underlying cell-intrinsic effects, we trained another model by regressing features on intrinsic age, while accounting for mean number of counts per subject. Genes were sorted by the difference in the intrinsic age coefficient and age coefficient to extract genes more strongly associated with intrinsic age relative to chronological age in the negative or positive direction. Gene lists were passed to gseGO() function with minGSSize = 3, maxGSSize = 1000 and pvalueCutoff = 0.01. Gene annotations for homo sapiens were used (org.Hs.eg.db).

### Hypergeometric Test for Over- and Under-representation Analysis

Hypergeometric cumulative distribution function was applied to evaluate whether there’s a significant deviation in the proportion of a cell type or state in a particular group of subjects from the expected proportion in the total population (null hypothesis). This was applied in two contexts, first is calculating the likelihood of observing as few cells or less (lower tail of the distribution, underrepresentation) or as many cells or more (upper tail of the distribution, overrepresentation) of a particular cell type or cell state. In terms of cell type composition, we estimated the underrepresented cell type populations in old relative to young individuals ^20^ and AD patients relative to age-matched controls ^38^. To quantify the disparity in the abundance of cells depending on the enriched gene signature, overrepresentation of cells enriched for positively or negatively associated signatures with systemic and intrinsic age per cell type was estimated.

### Epigenomic Erosion Scoring Per Cell

We computed the epigenomic erosion score per cell based on ChromHMM states for the dorsolateral prefrontal cortex (E073) ^77^. Our workflow followed the previously described scoring method with minor adjustments ^46^: (1) Peak calling using cell type annotations in ArchR’s addGroupCoverages, addReproduciblePeakSet and addPeakMatrix functions, (2) Extraction of genomic coordinates for reproducible peaks, (3) Annotation of peaks with ChromHMM 18 states using bedtools intersect function, (4) Annotation of the binarized peak matrix for excitatory and inhibitory neurons based on the annotated reads, (5) For each cell, the number of counts per chromatin state by divided by the total number of counts for the remaining states to compute the fraction of reads per state resulting in *n cells*18 states* matrix, (6) Center-log-Normalization per state across cells, (7) Assigning state weights to each state value per cell, where Active states (Enhancers; TSS; Tx) assigned negative sign, and the repressive regions (Quies, Repr, Het, TxWk) assigned positive sign, (8) For each cell, an epigenomic score is calculated by summing up scores across the 18 states, higher scores (more positive) represent more opening of repressive regions and higher erosion.

### Differential Chromatin Accessibility Analysis for Human Prefrontal Cortex snRNA-Seq

Using the Limma-Voom approach ^78,79^, we performed a differential chromatin accessibility analysis to identify regulatory elements specific to inhibitory neurons enriched in the negative intrinsic age signature while accounting for donor-level variation across pseudobulk samples. This analysis consisted of the following steps: **(1) Cell subsetting:** snATAC-Seq data from the Anderson multiomics dataset were analyzed using ArchR. Analyses focused on inhibitory neurons from Alzheimer’s disease (AD) donors. **(2) Pseudobulk aggregation:** cells were grouped at the pseudobulk level by donor and intrinsic gene set state (binary variable: negative intrinsic vs. non-negative intrinsic), generating a grouping variable that allowed binary gene set-specific profiles per individual while retaining donor-level structure. Reproducible peaks were obtained from the ArchR project. We constructed pseudobulk peak accessibility matrices using getGroupSE, aggregating peak counts across cells within each donor × gene set group. **(3) Peak filtering:** To enrich for regulatory elements, peaks annotated as distal or intronic and located more than a 10 kb from transcription start sites (to exclude promoters) were retained. **(4) Model Design:** we constructed a design matrix model (∼ 0 + Intrinsic Geneset State + nCells + ReadsInPeaks + FRIP), capturing intrinsic age signature enrichment effects on chromatin accessibility while controlling for technical covariates (number of cells, total reads/fragments in peaks and fraction of reads in peaks per pseudobulk sample). **(5) Normalization and Variance modeling:** Peaks with low average accessibility were excluded prior to modeling. Library size normalization factors were computed using calcNormFactors(). We estimated observation-level precision weights modeling the mean-variance relationship and sample-specific quality weights using voomWithQualityWeights(). To account for repeated measurements from the same subject, we computed intra-subject correlation using duplicateCorrelation() function, then recomputed voom precision weights incorporating sample correlations. **(6) Linear Modeling and Contrasting:** we fitted a linear model with lmFit() and applied empirical Bayes moderation to stabilize variance estimates with eBayes(). Gene set-specific contrasts were defined to compare intrinsic negative aging states against all other states. **(7) Covariate-Adaptive False Discovery Weight:** For the differentially accessible peaks, we applied multiple hypothesis correction using the FDR method while accounting for average accessibility as a covariate using lm_qvalue() from swfdr package ^80^. Peaks were considered significantly accessible if they met stringent statistical thresholds: peaks were first ranked by effect size, and those in the top 2% in the positive direction (≥ 98th percentile) were retained. This was followed by adjusting significance and filtering peaks with adjusted FDR < 0.05.

### Transcription Factor Motif Enrichment

Motif enrichment analysis was performed using HOMER on sets of differentially accessible peaks, with the hg38 genome and default background settings. Known motif results were filtered for highly significant enrichments and processed to extract motif identity and enrichment strength. Motif enrichment effect size is reported as the difference in the percentage of target peaks versus background regions containing each motif (Δ% = % targets − % background), reflecting how specifically each motif is enriched above the genomic background.

### Evolutionary Conservation Analysis of Enhancer-Gene Synteny Across Vertebrates

To evaluate evolutionary conservation of enhancer-gene relationships, human enhancer coordinates were projected across vertebrate genomes using the 577-species Vertebrate Genomes Project (VGP) whole-genome alignment. To identify enhancer-gene relationships, each enhancer was assigned to its nearest gene based on transcription start site (TSS) proximity. In the human reference genome (hg38), the nearest TSS was identified for each enhancer using BEDTools. Human enhancer and TSS coordinates were then projected to each vertebrate genome using HAL Liftover ^72^, and the nearest lifted TSS was identified for every lifted enhancer. These nearest-gene assignments were used to assess conservation of enhancer-gene synteny across species. Enhancers lacking a lifted coordinate or a corresponding TSS annotation in a given species were excluded from downstream analyses.

To quantify conservation of enhancer-gene synteny, the nearest gene associated with each enhancer in a target species was compared with the nearest gene assigned to the orthologous enhancer in hg38. Enhancers were classified as synteny-retained when the nearest-gene assignment matched the human reference and as non-retained otherwise. For each enhancer, we additionally calculated the change in enhancer-gene distance relative to hg38 by subtracting the human enhancer-to-TSS distance from the corresponding distance in the target species. Species-level summary statistics were calculated by aggregating enhancer measurements within each genome. These included the percentage of enhancers retaining synteny, the median absolute enhancer-to-TSS distance, the median enhancer-gene distance shift relative to hg38, and the proportion of enhancers successfully lifted. Genome metadata, including taxonomic classification and divergence time estimates, were obtained from the Vertebrate Genomes Project metadata table. Species were grouped into major vertebrate clades (mammals, birds, reptiles, amphibians, and fish), and divergence times from humans were used for comparative analyses.

To determine whether aging-vulnerable enhancers exhibited distinct evolutionary properties, identical analyses were performed on a size-matched set of randomly selected background enhancers. Species-level statistics from vulnerable and random enhancer sets were compared across vertebrates. The relationship between enhancer-gene distance shifts and evolutionary divergence was evaluated using linear regression models, with log-transformed median absolute distance or distance shifts modeled as a function of divergence time (millions of years since divergence from the human lineage), enhancer class (vulnerable versus random), and their interaction: ***lm(log10(median distance + 1) ∼ divergence time × enhancer class)***.

## CODE AND DATA AVAILABILITY

Datasets used for modeling and downstream analyses are publicly available ^9,20,38,40,52^. The iENR framework is implemented using elastic net regression with iterative feedback-based feature selection. All R and Python scripts used for data processing, analysis and generation of main and Extended Data Figures are available at: https://github.com/pfenninglab/brain-aging-evolution/. Any additional custom code or resources not included in the repository can be requested from the corresponding author.

## AUTHOR CONTRIBUTIONS

G.A. contributed to hypothesis generation, computational analysis design and implementation, data processing, iterative aging model design, training, optimization, and evaluation, as well as manuscript writing and editing and figure generation. Q.S. contributed to data processing and preparation for model training and trained earlier versions of the aging clock models. A.Z.W. contributed to data processing and preparation for model training, as well as the training, optimization, and evaluation of cortical TACIT models. R.G. contributed to synteny analysis across VGP genomes. B.N.P. contributed to data processing and provided guidance on the cortical TACIT model. H.H.S. contributed to the generation of controlled versions of the cortical TACIT pipeline. V.C. contributed to differential accessibility analysis. The VGP consortium provided high-quality data for 577 vertebrate genomes. A.R.P. provided the conceptual design of the aging model, guided the analytical strategy, and provided critical feedback on the analyses and manuscript.

## ACKNOWLEDGMENTS

We would like to thank Irene Kaplow, Michael Kleyman, Michael Gee, Aaron Lewis and Nikhil Yadala for their contributions to earlier versions of the TACIT and molecular aging clock models. We like to acknowledge funding from the Michael J. Fox Foundation (ASAP-025193) and National Institute of Health (NIH NIA R01 AG077636-01).

